# RNA Methylome Reveals the m^6^A-Mediated Regulation of Flavor Metabolites in Tea Leaves under Solar-Withering

**DOI:** 10.1101/2022.05.12.491608

**Authors:** Chen Zhu, Shuting Zhang, Chengzhe Zhou, Caiyun Tian, Biying Shi, Kai Xu, Linjie Huang, Yun Sun, Yuling Lin, Zhongxiong Lai, Yuqiong Guo

**Author notes:** Corresponding authors. (Guo Y), (Lai Z).

## Abstract

Epitranscriptomic mark *N*^6^-methyladenosine (m^6^A) is the most predominant internal modification in RNAs, which plays pivotal roles in response to diverse stresses. Multiple environmental stresses caused by withering process can greatly influence the accumulation of specialized metabolites and the formation of tea flavor. However, little is known about the effects of m^6^A-mediated regulatory mechanism on flavor-related metabolisms in tea leaves. Here, we explored m^6^A-mediated regulatory mechanism and its impacts on flavonoid and terpenoid metabolisms under solar-withering using integrated RNA methylome and transcriptome. Dynamic changes in global m^6^A levels of tea leaves are mainly controlled by two m^6^A erasers (CsALKBH4A and CsALKBH4B) under solar-withering. Differentially methylated peak (DMP)-associated genes under different shading rates of solar-withering were identified and found to be enriched in terpenoid biosynthesis and spliceosome pathways. Further analyses indicated that CsALKBH4-driven RNA demethylation can not only directly affect the accumulation of volatile terpenoids by mediating the stability and abundance of terpenoid biosynthesis-related genes, but also indirectly influence the contents of flavonoids, catechins, and theaflavins via triggering the alternative splicing (AS)-mediated regulation. Our findings underscored a novel layer of epitranscriptomic gene regulation in tea flavor-related metabolic pathways and established a compelling link between m^6^A-mediated regulatory mechanism and the formation of high-quality flavor in tea leaves under solar-withering.

## Introduction

As the most predominant internal modification in eukaryotic mRNAs, *N*^6^-methyladenosine (m^6^A) is recognized as a critical transcriptional regulator that establishes a novel molecular mechanism for profoundly affecting various biological processes via modulating multiple aspects of mRNA processing and metabolism [1], including mRNA abundance [2,3], stabilization [4], and splicing [5]. In analogy to well-studied epigenetic regulation mediated by modifications to DNAs and histone proteins, the dynamic and reversible m^6^A modification is dominantly installed and removed by m^6^A methyltransferases (writers) and demethylases (erasers), respectively. In addition to these two crucial components, m^6^A readers, are responsible for the localization and recognition of m^6^A-containing RNAs to implement the biological effects of m^6^A modification. Consequently, m^6^A writers, erasers, and readers collaboratively orchestrate a complicated regulatory network that governs the m^6^A modification. Although great progress has been achieved in understanding the potential functions of m^6^A modification in animals, the current study devoted to m^6^A regulatory mechanism in plant kingdom is just beginning to be disclosed. Reportedly, m^6^A-mediated regulatory mechanism has been shown to affect the normal growth of root and shoot in *Arabidopsis thaliana* [6,7]. Furthermore, an increasing number of studies suggest that m^6^A modification is also involved in the regulating response to diverse environmental stresses, including drought [8], cold [9], and ultraviolet (UV) radiation stresses [10]. More recently, a few studies have started on the precise functions of m^6^A regulatory mechanism or m^6^A regulatory genes in plant kingdom [11–14]. Compared with model plants, regarding the regulatory mechanisms as well as the functional roles of m^6^A modification, the evidence directly from investigations in horticultural crops is largely limited.

Tea (*Camellia sinensis*) plant is an important and traditional economic crop and is cultivated on a massive scale in many developing and developed counties. Its tender buds and leaves are mostly used to produce the most consumed and popular beverage for millennia. Oolong tea, one of the six tea categories in China, is famous for its elegant floral and fruity aroma, as well as unique brisk-smooth and mellow taste. The flavory and sensory characteristics of oolong tea largely depend on the postharvest manufacturing process [15]. During manufacturing of oolong tea, withering is the first essential stage that impacts palatability and commercial value of tea. The picked leaves are still in a live state and inevitably encounter various environmental stresses during tea-withering stage, concomitant with obvious alterations in multiple taste and aroma compounds, as well as endogenous phytohormones, which directly or indirectly endows oolong tea with characteristic flavor and health benefits [16–18]. Among these taste compounds, flavonoids are a large group of secondary metabolites in tea plants and are considered to be tightly related to the tea palatability, especially catechins, which are the most dominant component of flavonoids and comprise 12%–24% of tea dry weight [19]. It is generally accepted that flavonoids and catechins are the major contributors to the velvet astringency and bitterness of tea [20]. Moreover, catechins can be further oxidized to the high-molecular-weight polymeric compounds referred to as theaflavins, which confer tea with beneficial health-promoting properties and characteristic mellow taste [21]. As the most representative aroma compounds in oolong tea, volatile terpenoids are the cornerstone of high-quality oolong tea because of their contributions to the pleasant floral and fruity fragrance. Decades of studies have demonstrated that multiple environmental stresses caused by withering treatments result in a series of multidimensional responses extending from genes to metabolites, and further affects the formation of oolong tea flavor [22,23]. However, our knowledge regarding the upstream regulation mechanism of the flavor-related genes and relevant metabolites is still limited.

In the previous study, we uncovered that solar-withering is a conventional withering method in tea production, which is beneficial to the development of high-quality flavor in oolong tea [24,25]. Light, as an energy source and signal molecule, provides the prerequisite conditions for metabolic changes under solar-withering. Typically, light quality and intensity are crucial parameters that determine the effects of solar-withering on the tea quality. Prior studies have largely focused on the mechanism underlying the effects of light quality on the tea flavor [26,27]. Hitherto, few researchers have hammered at the impacts of different shading rates on the metabolism of tea flavor compounds and formation of tea flavor during solar-withering stage. In actual production, the establishment of traditional solar-withering method relies heavily on the subjective experiences and random trials of tea makers, leading to significant instability in the tea quality. To overcome this limitation, understanding the metabolic pathways that affect flavor formation, and their potential regulatory mechanisms under different shading rates of solar-withering, is the pivotal to optimizing the utilization of the shading rate and the efficient production of high-quality oolong tea. More recently, accumulating evidence has indicated that epigenetic factors, including DNA methylation and histone modifications, portray crucial roles in the biosynthesis of secondary metabolite and tea aroma formation [28,29]. However, the functional roles of RNA methylation, another epigenetic modification, and precise regulatory mechanisms underlying the m^6^A-mediated flavor formation in the tea-withering stage remain largely unanswered. Fortunately, successful decoding of tea chromosome-scale genome has laid a solid foundation for the investigation of m^6^A modification in tea plant [30].

In the present study, integrated RNA methylome and transcriptome analyses were applied to investigate the effects of m^6^A modification on the flavor formation in tea leaves under different shading rates of solar-withering. We systematically identified the differentially methylated peak (DMP)-associated genes, and revealed that the alteration of m^6^A modification in many DMP-associated genes is inextricably associated with the solar-withering treatment, especially the shading rate of withering. We further demonstrate that, CsALKBH4*-*mediated RNA demethylation alters the m^6^A abundance within 3′UTR and near stop codon and regulates the expression levels of m^6^A-modified RNAs, thereby affecting the accumulation of flavor metabolites and tea palatability. The roles of m^6^A-mediated alternative splicing (AS) regulatory mechanism under solar-withering were also explored. These findings generated herein are a crucial first step towards understanding the regulatory effects of m^6^A modification on tea plant and provide deeper insight into the roles of m^6^A-mediated regulatory mechanism in the development of high-quality oolong tea under solar-withering.

## Results

### Significant change in the phenotypes and global m^6^A abundance of tea leaves under solar-withering

The light intensity, spectrum, and UV intensity of solar-withering with different shading rates were monitored (Figure S1). The detailed withering treatments were performed as follows: solar-withering with high shading rate (SW1); solar-withering with middle shading rate (SW2); solar-withering with low shading rate (SW3); solar-withering with natural sunlight (SW4). Notably, the light intensity and UV intensity under the sunshade net decreased coordinately with the elevation of shading rate. However, the utilization of sunshade nets has little effect on spectral composition, light quality, and wavelength compared with the natural sunlight. Significant differences in UV intensity were found between solar-withering with and without shading net. The shading treatment greatly reduced the UV intensity. From SW1 to SW3, different shading rates have relatively little effect on the UV intensity. Next, the external phenotypes of fresh leaves (FL) and four solar-withered leaves with different shading rates were recorded (**Figure 1A**). It was obvious that the fresh leaves were the straight, glossy, and jade green. With the decrease in the shading rate of solar-withering, the withered leaves become more shrunken and deformed, and leaf color gradually darkened. We then evaluated the global m^6^A/A ratio of these tea leaves (Figure 1B). Solar-withering treatment repressed global m6A abundance in four solar-withered leaves with different shading rates. The overall m^6^A level gradually decreased from FL to SW3, while the m^6^A abundance exhibited an obvious rebound at SW4.

**Figure 1.**
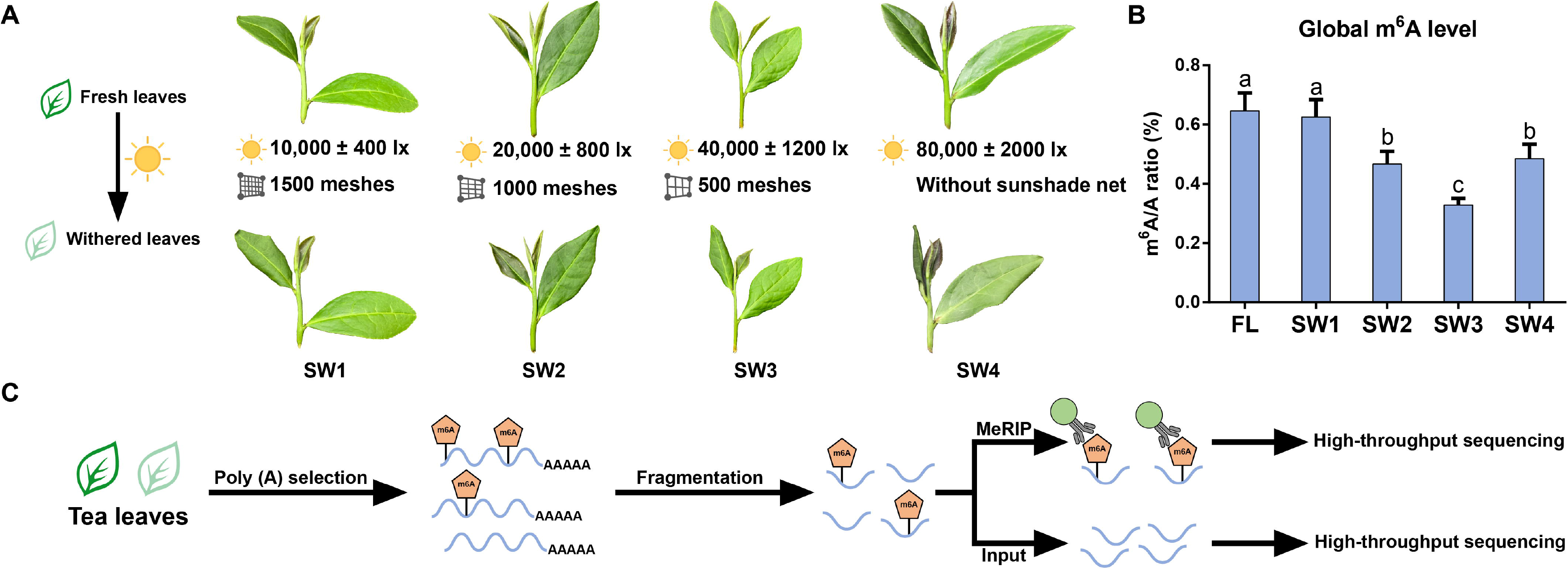
Flowchart of the study on m^6^A methylation in tea leaves under solar-withering. **A.** The external phenotypes of withered leaves before and after solar-withering with different shading rates. **B.** Dynamics of global m^6^A levels in tea leaves under solar-withering with different shading rates. Data are presented as mean ± standard deviation. Different letters indicate a significant difference (*P* < 0.05). **C.** Brief workflow for MeRIP-seq. The fragmented RNA was divided into two portions, one of which was enriched with m^6^A-specific antibody in IP buffer. The other RNA without IP treatment was used as the input control. Finally, the two portions of RNA were sequenced using an Illumina NovaSeq 6000 platform. FL, fresh leaves; SW1, solar-withering with high shading rate; SW2, solar-withering with middle shading rate; SW3, solar-withering with low shading rate; SW4, solar-withering with natural sunlight; MeRIP-seq, methylated RNA immunoprecipitation sequencing; IP, immunoprecipitated.

### Overview of m^6^A MeRIP-seq

To ascertain the functional roles of m^6^A-mediated regulatory mechanisms under solar-withering, we performed m^6^A MeRIP-seq to analyze m^6^A modification at the mRNA level in tea leaves treated with different shading rates of solar-withering (Figure 1C). A total of 34–50 and 63–76 million high-quality clean reads were obtained for immunoprecipitated (IP) libraries and input libraries, respectively, among which 28–41 and 55–67 million reads were accordingly mapped to tea genome (Table S1). The Q20 and Q30 indexes of each library exceeded 93% and 88%, respectively. Only the m^6^A peak in all biological replicates showing the high-confidence coefficients was selected for further analysis. Overall, a total of 3570, 3249, 2966, 2809, and 2883 high-confidence m^6^A peaks were identified from FL, SW1, SW2, SW3, and SW4, respectively (Figure S2). Among these peaks, 4605 m^6^A peaks were newly generated after solar-withering treatment. Additionally, we identified a total of 31,720, 31,958, 32,146, 32,215, and 31,988 mRNA transcripts were obtained from fresh leaves and four solar-withered leaves with different shading rates, respectively. Next, we identified 5277 high-confidence m^6^A peaks originating from 4289 mRNA transcripts. Obviously, 1988 m^6^A-marked genes were present in all samples (Figure S3). Solar-withering treatment induced the production of 843 new m6A-marked genes. The variation trend in the number of m6A-marked genes under solar-withering with different shading rates were in line with the dynamics of total m^6^A abundance. These data suggest that m^6^A abundance was closely related to these changes in the number of m6A-marked genes, similar to previous findings in tomato [11]. The amount of m^6^A-marked genes gradually decreased from FL to SW3, whereas the number of m^6^A-marked genes increased in the SW4 compared with SW3. Besides, most of the m^6^A-marked genes (2229 genes) were common in FL *vs*. SW2 and FL *vs*. SW4 comparisons, whereas 219 and 216 m^6^A-marked genes were uniquely detected in the FL *vs*. SW2 and FL *vs*. SW4 comparisons, respectively. These results suggest that m^6^A-marked genes were not significantly different in these two comparisons, and most m^6^A-marked genes were shared in these comparisons. Of the 4289 m^6^A-marked genes, most genes in the five samples (an average of 96.90%) contained a single m^6^A peak, while only a few transcripts harbored more than three m^6^A peaks (Table S2). The distribution of m^6^A peaks in the transcriptome-wide of tea plants was then investigated. Major m^6^A modifications within mRNA transcripts were primarily positioned in the adjacent of stop codon and 3′UTR (**Figure 2A** and B), while only 1.46%–2.13% of m^6^A enrichment was located in CDS, 5′UTR, and neighboring start codon (Figure 2B). From FL to SW3, the proportion of m^6^A modifications in 3′UTR and near stop codon increased, while the proportion located in 5′UTR, CDS, and near start codon decreased. Surprisingly, a decrease in the proportion of m^6^A enrichment was observed in 3′UTR and around stop codon in SW4 compared to that in SW3, but the proportion of m^6^A peaks within 5′UTR, CDS, and neighboring start codon increased in SW4. Next, the m^6^A peaks were normalized by enrichment algorithm [31], and the results revealed that m^6^A modifications were accumulated preferentially in 3′UTR and around stop codon (Figure S4), which was analogous to the previous description of several plant species [12,32,33]. Overall, the distribution of m^6^A peak in tea leaves under solar-withering changed slightly to some extent. This is in accordance with that observed in tomato, which did not display apparent variations in distribution of m^6^A modifications during fruit ripening [11].

**Figure 2.**
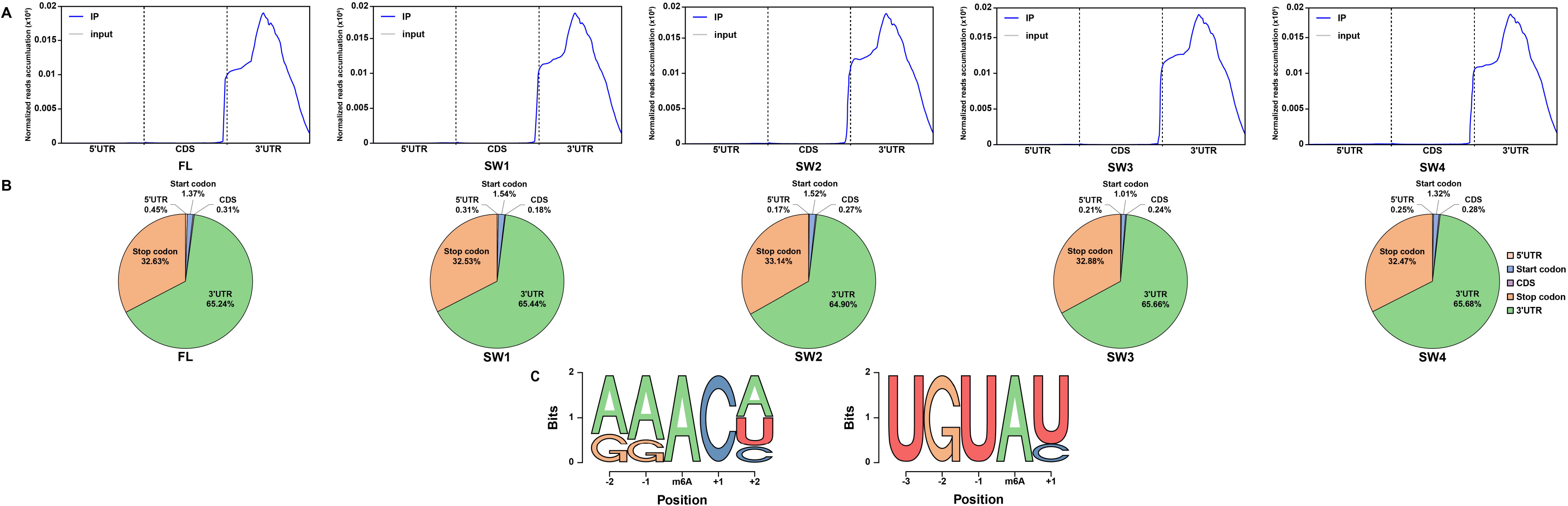
Characteristics of m^6^A distribution and sequence motifs in tea leaves under solar-withering with different shading rates. **A.** Metagenomic profiles of m^6^A distribution along transcripts. **B.** Pie charts showing the fraction of m^6^A peaks fell into five transcript segments (5′UTR, start codon, CDS, stop codon, and 3′UTR). **C.** Top sequence motifs identified within m^6^A peaks. FL, fresh leaves; SW1, solar-withering with high shading rate; SW2, solar-withering with middle shading rate; SW3, solar-withering with low shading rate; SW4, solar-withering with natural sunlight; 5′UTR, 5′ untranslated region; CDS, coding sequence; 3′UTR, 3′ untranslated region.

To obtain the enriched motifs within the confines of m^6^A peaks in tea plants, all m^6^A peaks were scanned using MEME suite and HOMER tool. As expected, we observed that the canonical motif RRACH (R=G or A, H=A, U, or C) is noteworthy enriched within the most of m^6^A peaks (Figure 2C). Besides, we identify another enriched motif UGUAY (Y=C or U), which was similar to a plant-specific motif URUAY that can be recognized by m^6^A readers [34]. This motif UGUAY is primarily observed in m^6^A peaks of tomato [11], but not detected in other plant species. This result showed that the enriched motif of m^6^A-modified sites was relatively conserved in two horticulture plants, tea plant and tomato, compared to other plant species.

### Analysis of DMP-associated genes

To gain an insight to the potential effect of m^6^A methylation under solar-withering, we first searched for DMPs in the entire m^6^A methylome. DMPs were identified according to the criteria of |fold change (FC)| ≥ 2 and *P* < 0.05. A total of 265 DMPs were detected in the FL *vs*. SW1 comparison, comprising 203 hypermethylated peaks and 62 hypomethylated peaks (Table S3). Besides, 215 DMPs were found in the FL *vs*. SW4 comparison, which was close to 217 DMPs detected in the FL *vs*. SW2 comparison and more than 183 DMPs identified in the FL *vs*. SW3 comparison. These data showed that apparent changes in the global m^6^A status occurred at withered leaves under solar-withering with different shading rates. The accumulation of detected DMPs continued to decreased from FL *vs.* SW1 to FL *vs.* SW3 comparisons, whereas the number of DMPs was dramatically elevated in the FL *vs.* SW4 comparison. Intriguingly, we observed that the change in the number of DMPs under solar-withering was in accordance with the dynamic trend of total m^6^A level. These results suggested that the alteration of m^6^A modification in tea leaves was tightly associated with the solar-withering treatment, especially the shading rate of tea-withering.

Evidence is accumulating that m^6^A deposition affect the mRNA levels under different developmental stages or environmental stresses [9,13,32,35]. To evaluate the overall correlation between m^6^A modification and gene expression levels under solar-withering, a total of 1137 DMP-associated genes with altered expression levels between samples were selected (Figure S5). The results showed that 74.9% of DMP-associated genes in FL *vs*. SW1 comparison, 79.6% of DMP-associated genes in FL *vs*. SW2 comparison, 83.2% of DMP-associated genes in FL *vs*. SW3 comparison, and 79.6% of DMP-associated genes in FL *vs*. SW4 comparison with increased or decreased m^6^A enrichment negatively regulated the gene expression, whereas only 25.1% in FL *vs*. SW1 comparison, 20.4% in FL *vs*. SW2 comparison, 16.8% in FL *vs*. SW3 comparison, and 20.4% in FL *vs*. SW4 comparison with increased or decreased m^6^A enrichment positively influenced the gene expression. These data implied that m^6^A enrichment in most DMP-associated genes was negatively correlated with gene expression levels under tea-withering. Intriguingly, we noticed that, combined with the elevated proportion of m^6^A modifications in 3′UTR and near stop codon from FL to SW3, m^6^A modification within these regions tended to negatively affect the mRNA abundance. Besides, the proportion of m^6^A modification within 3′UTR and around stop codons was reduced in SW4 compared to SW3, concomitant with an obvious reduction in the proportion of DMP-associated genes showing a negative correlation between m^6^A abundance and expression levels. Concretely, m6A peaks distributed within 3′UTR and around stop codon showed a positive correlation with the mRNA abundance, whereas m6A deposition located in 5′UTR, CDS, and near start codon tended to be negatively correlated with the gene expression. These findings implied that the multifaceted effects of m^6^A modification on mRNA expression may depend on the position of m^6^A peaks in gene structure. The specific feature of m^6^A distribution in tea leaves is closely related to gene expression, and it is conserved among several important crops, including rice [36], strawberry [13], and apple [37].

Kyoto encyclopedia of genes and genomes (KEGG) enrichment analysis was implemented to decipher the biological functions of all identified DMP-associated genes. In line with the effects of tea-withering on flavor formation, a large number of m^6^A-labeled genes were found to be remarkably enriched in KEGG functional class associated with genetic information processing (**Figure 3A**). More than half of DMP-associated genes were primarily enriched in ribosome, RNA transport, and spliceosome pathways. Besides, we found that several DMP-associated genes were annotated to three KEGG pathways belonging to another functional class “metabolism”. Among these pathways, terpenoid backbone biosynthesis pathway contained the highest value of rich factor, and it was closely related to the aroma formation of oolong tea. We therefore spotlighted on these four KEGG pathway in further analyses.

**Figure 3.**
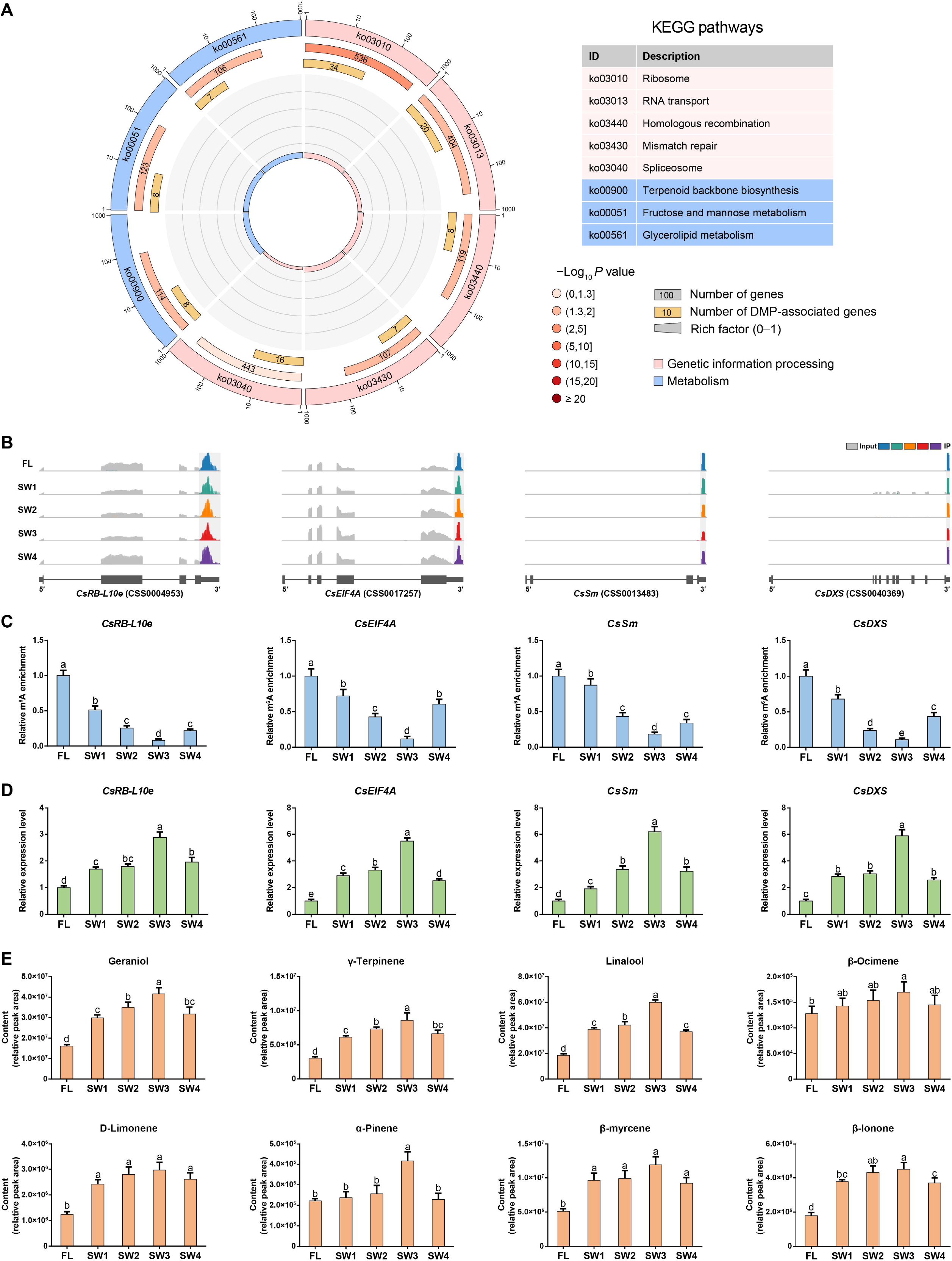
m^6^A abundance of representative DMP-associated genes was negatively correlated with their expression levels and the accumulation of volatile terpenoids under solar-withering with different shading rates. **A.** KEGG enrichment analysis of DMP-associated genes. From the outside to the inside, the first circle indicates enriched KEGG pathways and the number of the genes corresponds to the outer circle. The second circle indicates the number of the genes in the genome background and *P* value for enrichment of the genes for the specified KEGG pathway. The third circle indicates the number of the DMP-associated genes. The fourth circle indicates the enrichment factor of each KEGG pathway. **B.** The distribution of m^6^A reads in *CsRB-L10e*, *CsEIF4A*, *CsSm*, and *CsDXS*. Exons and introns in the gene structures are represented by the thick boxes and lines, respectively. **C.** Relative m^6^A enrichment of *CsRB-L10e*, *CsEIF4A*, *CsSm*, and *CsDXS* under solar-withering with different shading rates determined by m^6^A-IP-qPCR. **D.** Relative expression levels of *CsRB-L10e*, *CsEIF4A*, *CsSm*, and *CsDXS* under solar-withering with different shading rates determined by qRT-PCR. **E.** Relative contents of volatile terpenoids under solar-withering with different shading rates determined by GC-MS. Data are presented as mean ± standard deviation. Different letters indicate a significant difference (*P* < 0.05). FL, fresh leaves; SW1, solar-withering with high shading rate; SW2, solar-withering with middle shading rate; SW3, solar-withering with low shading rate; SW4, solar-withering with natural sunlight; DMP-associated genes, differentially methylated peak-associated genes; KEGG, Kyoto encyclopedia of genes and genomes; m^6^A-IP-qPCR, m^6^A-immunoprecipitation-qPCR; qRT-PCR, quantitative real-time polymerase chain reaction; GC-MS, gas chromatography-mass spectrometry.

### A negative correlation between m^6^A abundance and expression levels in most DEP-associated genes under solar-withering

In the m^6^A-seq analysis, we observed that the m^6^A abundance and expression levels of several core genes involved in four aforementioned KEGG pathways were significantly altered during tea-withering process (Figure S6). In ribosome pathway, a total of 34 ribosomal protein-encoding genes (RP-genes) exhibited differential m^6^A abundance under solar-withering (Table S4). These RP-genes are indispensable in ribosome formation and protein biosynthesis, and play a significant role in the transcription-translation process. Besides, we observed that 20 and 16 DMP-associated genes were assigned to RNA transport and spliceosome pathways, respectively (Table S5 and S6). RNA transport is functionally linked to several steps in RNA processing stage, including RNA splicing, 3′-end formation, and transcription-translation processes, which is the fundamental for genes to perform various biological functions [38]. Concomitantly, AS is a pivotal post-transcriptional process to enhance diversities of transcripts and corresponding proteins, and is a critical molecular mechanism for regulating the transcriptional abundance of coding genes in the plant kingdom. Interestingly, eight terpenoid biosynthesis-related genes showed significant changes at m^6^A levels during tea-withering stage (Table S7). It is well-known that the floral and fruity aroma of oolong tea is mainly attributed to its characteristic terpenoids [17]. Thus, we indicated that RNA methylation may affect the tea flavor by regulating the m^6^A modification of terpenoid biosynthesis-related genes.

Among the four above-mentioned pathways, we selected one DMP-associated gene in each pathway that exhibited the dynamic alteration at the m^6^A level under solar-withering with different shading rates. These four genes, including *large subunit ribosomal protein L10e-encoding gene* (*RB*-*L10e*), *eukaryotic translation initiation factor 4A* (*EIF4A*), *core spliceosomal Sm protein-encoding gene* (*Sm*), and *1*-*deoxy*-*D*-*xylulose*-*5*-*phosphate synthase gene* (*DXS*), were warrant further investigation. The m^6^A peaks were mainly distributed within 3′UTR and around stop codon in mRNA structure of the selected genes (Figure 3B), and significant changes in the m^6^A levels of these four genes were observed in our m^6^A-seq datasets (Figure S7A). We then performed the m^6^A-immunoprecipitation-qPCR (m^6^A-IP-qPCR) to further validate the m^6^A enrichment of four above-mentioned genes retrieved from m^6^A-seq data. As expected, the m^6^A abundance of *CsRB-L10e*, *CsEIF4A*, *CsSm*, and *CsDXS* dropped sharply from FL to SW3, but exhibited a distinct rebound from SW3 to SW4 (Figure 3C). In comparison, these four genes showed sustained upward expression trends from FL to SW3, while the transcript levels of these genes were noticeably repressed in SW4, as revealed by both transcriptome datasets (Figure S7B) and quantitative real-time polymerase chain reaction (qRT-PCR) (Figure 3D). The results indicate that the m^6^A depositions of these mRNAs were negatively correlated with their corresponding expression levels, similar to a previous study that mRNAs with low m^6^A abundance in tomato tend to display higher expression [11]. Besides, *DXS* is thought to an indispensable gene in plastidial methylerythritol phosphate (MEP) pathway, a branch of terpenoid biosynthesis, and directs metabolic flux toward terpenoid biosynthesis to define the size of monoterpenoid and apocarotenoid pools [39]. In this study, volatile terpenoids were monitored under different shading rates of solar-withering. It is noticeable that obvious changes in the contents of seven monoterpenoids and one apocarotenoid (Figure 3E). The abundance of all these eight volatiles increased markedly from FL to SW3 and reached a peak at SW3, but showed a significant decline at SW4. Concomitantly, we observed a positive relationship between the accumulation of eight volatile terpenoids and the *CsDXS* expression, while the abundance of these terpenoids was negatively correlated with m^6^A enrichment of *CsDXS* and overall m^6^A methylation level under solar-withering. These results implied that RNA methylation may affect the expression levels of terpenoid biosynthesis-related genes and terpenoid accumulation by modulating the m^6^A abundance of their respective genes.

### The expression profiles of m^6^A regulatory genes under solar-withering

The obvious alterations in m^6^A abundance were observed in a large number of gene related to involved in ribosome, RNA transport, spliceosome, and terpenoid biosynthesis pathways. Nevertheless, the roles of m^6^A-mediated regulatory mechanism in these pathways during tea-withering is still unclear. Reportedly, the function of RNA methylation is tightly associated with the transcript levels of RNA methyltransferase, demethylase, and reader genes [40]. Thus, we speculate that the m^6^A levels of these mRNAs are coordinately governed by m^6^A regulatory genes. According to our recent report [14], a total of 34 m^6^A regulatory genes were eventually identified from the chromosome-scale genome of tea plant. We initially analyzed the expression profiles of m^6^A regulatory genes under solar-withering (**Figure 4A**). The m^6^A regulatory genes with |FC| ≥ 2 and *P* < 0.05 were considered as differentially expressed genes (DEGs) between samples. The transcript levels of all m^6^A writers and readers were not altered obviously under solar-withering. Among 16 m^6^A eraser genes, only two m^6^A eraser genes, *CsALKBH4A* and *CsALKBH4B*, showed significant changes at the transcript level after solar-withering treatment. We also noticed that *CsALKBH6* transcription was only significantly induced under solar-withering with low shading rate (SW3). These results indicate that dynamic changes in m^6^A modification may be mainly controlled by m^6^A eraser genes, especially *CsALKBH6* played an important role in the regulation of m^6^A abundance in SW3. Next, qRT-PCR confirmed the expression patterns of m^6^A regulatory genes retrieved from transcriptome data and demonstrated that the expression levels of *CsALKBH4A* and *CsALKBH4B* increased continuously from FL to SW3 and peaked at SW3, but exhibited a distinct decrease at SW4 (Figure 4B). This trend was in accordance with the expression patterns of *CsRB-L10e*, *CsEIF4A*, *CsSm*, and *CsDXS*, but was inversely correlated with the m^6^A abundance of these four genes and global m^6^A level under solar-withering. Moreover, *CsALKBH6* expression was significantly elevated only in SW3, while m^6^A writer and reader genes were insensitive to solar-withering. These data implied that m^6^A eraser-mediated removal of m^6^A marks on mRNAs was closely linked with variations in m^6^A enrichment and expression abundance of DMP-associated genes.

**Figure 4.**
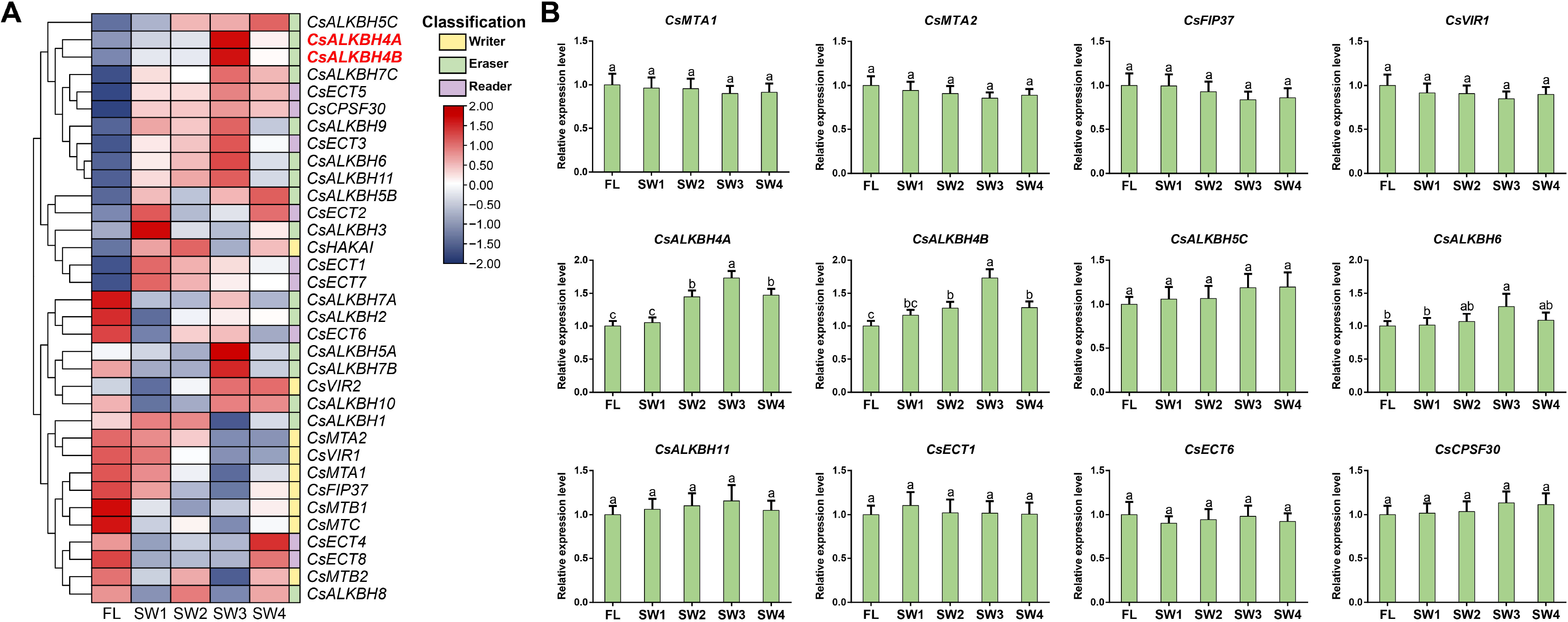
Expression profiles of m^6^A regulatory genes under solar-withering with different shading rates. **A.** The heatmap of m^6^A regulatory genes under solar-withering with different shading rates based on the transcriptome datasets. The expression values of m^6^A regulatory genes were normalized by log_2_-transformed (FPKM+1). **B.** Expression patterns of four m^6^A writer genes, five m^6^A eraser genes, and three m^6^A reader genes under solar-withering with different shading rates determined by qRT-PCR. Data are presented as mean ± standard deviation. Different letters indicate a significant difference (*P* < 0.05). FL, fresh leaves; SW1, solar-withering with high shading rate; SW2, solar-withering with middle shading rate; SW3, solar-withering with low shading rate; SW4, solar-withering with natural sunlight; FPKM, fragments per kilobase per million mapped reads; qRT-PCR, quantitative real-time polymerase chain reaction.

### CsALKBH4-mediated RNA demethylation may activate terpenoid biosynthesis

To gain further insights into the potential functions of *CsALKBH4A* and *CsALKBH4B* during tea-withering, we transiently suppressed the expression of these two genes via an siRNA-mediated gene silencing strategy (**Figure 5A**). As we anticipated, both *CsALKBH4A* and *CsALKBH4B* were significantly down-regulated in gene-silenced leaves, while the transcript levels of these two genes were not substantially altered by treatments with corresponding negative control (NC) (Figure 5B). Remarkably, the inhibitions of *CsALKBH4A* and *CsALKBH4B* led to a significant increase of overall m^6^A levels in gene-silenced tea leaves (Figure 5C), concomitant with an obvious elevation in m^6^A abundance of *CsRB-L10e*, *CsEIF4A*, *CsSm*, and *CsDXS* (Figure S8A), compared to the respective NC. Conversely, the mRNA levels of *CsRB-L10e*, *CsEIF4A*, *CsSm*, and *CsDXS* in gene-silenced leaves were markedly lower than those in NC-treated leaves (Figure S8B). These data hinted that the diminished expression of these four genes was mainly attributed to the suppression of CsALKBH4-mediated RNA demethylation. To further investigate how m^6^A demethylation influences the mRNA abundance of the *CsRB-L10e*, *CsEIF4A*, *CsSm*, and *CsDXS*, we next evaluated whether CsALKBH4-mediated RNA demethylation modulated mRNA stability of these four genes by monitoring the decay rate of mRNAs after actinomycin D treatment (Figure 5D). Accordingly, we observed that the mRNA abundance of *CsRB*-*L10e*, *CsEIF4A*, *CsSm*, and *CsDXS* degraded more rapidly in *CsALKBH4A*- and *CsALKBH4B*-silenced leaves compared with that in NC-treated leaves. It is notable that the decay rates of *CsRB*-*L10e*, *CsEIF4A*, *CsSm*, and *CsDXS* were positively correlated with their respective expression and negatively correlated with the m^6^A abundance of these four genes. These results suggest the excessive m^6^A sites in mRNAs of *CsRB*-*L10e*, *CsEIF4A*, *CsSm*, and *CsDXS* tend to destabilize these four mRNAs and result in an obvious plunge in transcript levels of these mRNAs. To assess the impacts of RNA demethylation on the accumulation of volatile terpenoids, the terpenoid contents in *CsALKBH4A-* and *CsALKBH4B*-silenced leaves were detected. The transcriptional repression of two *CsALKBH4* enormously decreased the amount of eight volatile terpenoids (Figure 5E). We speculate that, combined with the positive correlations between the abundance of eight volatile terpenoids and *CsDXS* expression, CsALKBH4-mediated RNA demethylation may directly activate terpenoid biosynthesis by removing m^6^A marks and enhancing the stability of corresponding mRNAs, thereby enormously improving the aroma quality of oolong tea under solar-withering.

**Figure 5.**
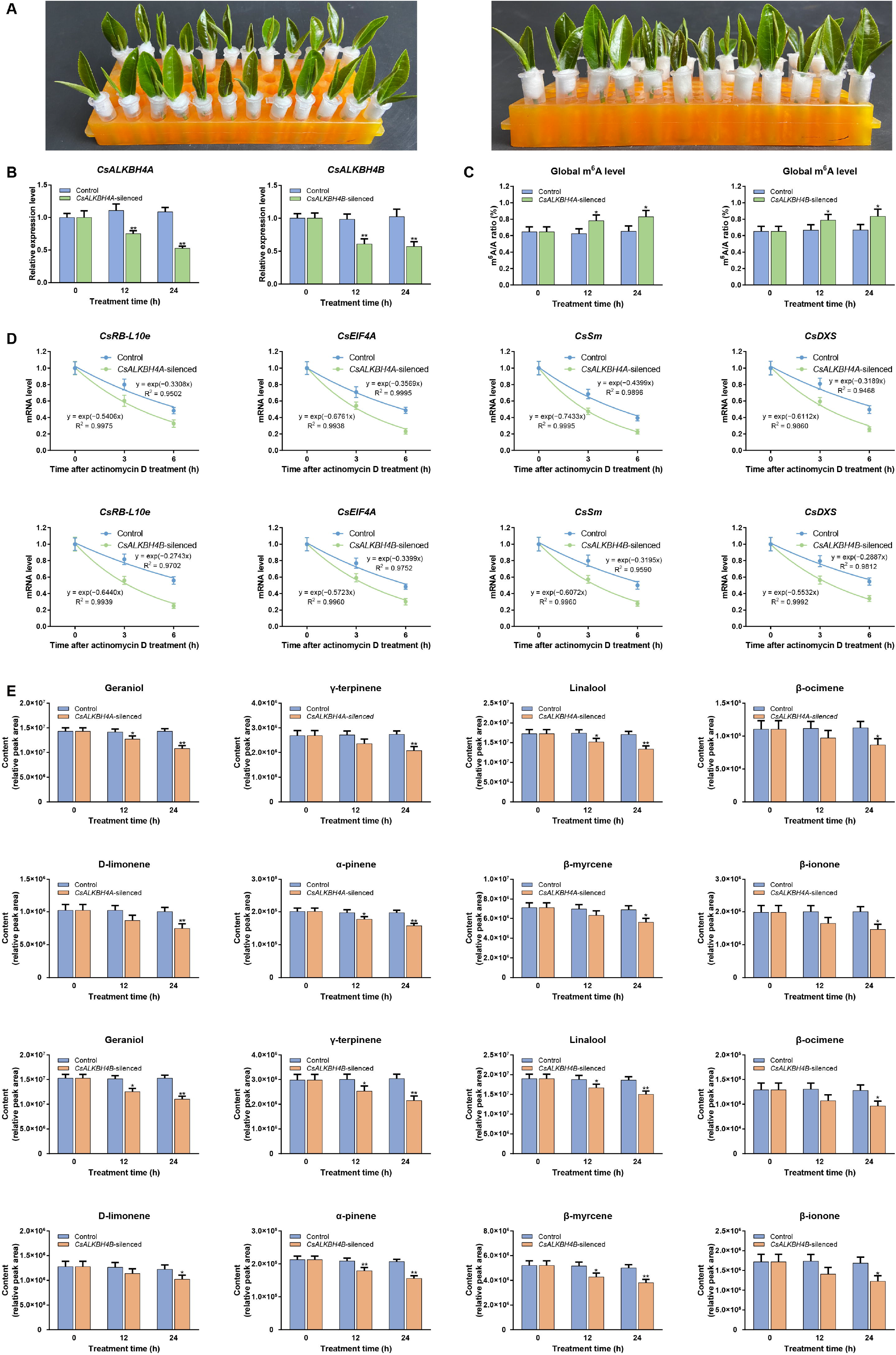
CsALKBH4-mediated RNA demethylation contributes to the accumulation of volatile terpenoids by removing the m^6^A marks and enhancing the stability of corresponding mRNAs. **A.** Diagram of *CsALKBH4A-* and *CsALKBH4B*-silenced assay in tea leaves via an siRNA-mediated gene silencing strategy. The freshly detached tea bud and first leaf from natural grown tea plants incubated in 1.5 mL microcentrifuge tubes that contained 1 mL of 20 μM siRNA or siRNA-NC solution. After incubation for 12 h and 24 h, the tea bud and first leaf were harvested and then used for qRT-PCR and metabolite detection. **B.** Expression levels of *CsALKBH4A* and *CsALKBH4B* in gene*-*silenced tea leaves determined by qRT-PCR. **C.** Dynamics of global m^6^A levels in gene*-*silenced tea leaves. **D.** mRNA stability of *CsRB-L10e*, *CsEIF4A*, *CsSm*, and *CsDXS* in gene*-*silenced tea leaves. To evaluate the mRNA stability, the leaf discs were collected from gene-silenced tea leaves and incubated in sterile water that contained 10 μg/mL actinomycin D solution. Tea leaves incubated in sterile water were as controls. Total RNAs were isolated from the leaves sampled at 6 h and 12 h, respectively. The mRNA levels of *CsRB-L10e*, *CsEIF4A*, *CsSm*, and *CsDXS* were examined by qRT-PCR. **E.** Relative contents of volatile terpenoids in gene*-*silenced tea leaves determined by GC-MS. Data are presented as mean ± standard deviation. Single asterisk indicates a significant difference (*P* < 0.05). Double asterisks indicate a highly significant difference (*P* < 0.01). qRT-PCR, quantitative real-time polymerase chain reaction; GC-MS, gas chromatography-mass spectrometry.

### RNA demethylation modulates AS events by influencing the m^6^A modification and expression of spliceosome-related genes

The preceding KEGG enrichment analysis of DMP-associated genes highlighted that the spliceosome pathway was one of the main KEGG enrichment pathways. The m^6^A modification of the core genes *Sm*, *pre-mRNA-splicing factor 18* (*Prp18*), and *Prp31* in the spliceosome pathway were significantly altered under solar-withering, concomitant with an obvious change in expression levels of these genes. Thus, we inferred that RNA demethylation might be implicated in the regulation of AS events by regulating the m^6^A abundance and expression levels of spliceosome-related genes. To assess this speculation, we firstly investigated the AS events under different shading rates of solar-withering. A total of 18,188 AS events were identified from 13,606 genes in five tea samples. After solar-withering treatment, the number of AS events in the solar-withered leaves were substantially increased relative to that in fresh leaves, suggesting the solar-withering may contribute to the occurrence of AS events (Table S8). Within a certain range of shading rate (from SW1 to SW3), the frequency of AS events was tightly related to the reduced shading rate of solar-withering. In contrast, solar-withering without the shading treatment led to a substantial decrease of AS events. Among these samples, retained intron (RI) were the predominant AS type. This observation is in accordance with that of previous reports on maize [41], cotton [42], and tea plant [43]. We next comprehensively identified the differentially expressed AS genes (DAGs) under different shading rates of solar-withering. The number of DAGs in FL *vs*. SW3 comparison was much greater than that in FL *vs*. SW1 and FL *vs*. SW2 comparisons, which indicated that the elevated intensity of solar-withering promoted the production of more DAGs (Table S9). Nevertheless, we noted that the number of DAGs in FL *vs*. SW4 comparison was fewer than that in FL *vs*. SW3 comparison. These data implied that solar-withering without the shading treatment attenuated the occurrence of DAGs during tea-withering stage. Meanwhile, the shading rate of solar-withering not only affected the alternation in the number of AS events, but also influenced the production of DAGs.

### The accumulation of flavonoids, catechins, and theaflavins was associated with the expression levels of AS transcripts under solar-withering

To further explore the putative effects of DAGs under solar-withering, we performed KEGG annotation on all identified DAGs. Most DAGs were enriched in functions related to metabolic pathways, followed by flavonoid biosynthesis pathway (**Figure 6A**). Therefore, structural genes involved in the flavonoid biosynthesis may be affected by the m^6^A-mediated AS regulatory mechanism (Figure S9, Table S10). We subsequently selected the two flavonoid biosynthesis-related DAGs. Of the two DAGs, *4*-*coumarate CoA ligase* (*4CL)* undergoing AS events generated two splicing variants, while another DAG *flavonoid 3’*-*hydroxylase* (*F3’H*) contained three AS transcripts. Based on the fragments per kilobase per million mapped reads (FPKM) value obtained from the transcriptome datasets, the expression profiles of these AS transcripts under solar-withering were analyzed (Figure 6B). The expression of full-length transcript *Cs4CL* substantially decreased from FL to SW3, but exhibited an obvious increase at SW4. The AS transcript *Cs4CL-a* had a similar expression profile to its full-length transcript. Concomitantly, the transcript abundance of *Cs4CL-a* was lower than that of *Cs4CL* under solar-withering stage. Solar-withering strongly also inhibited the transcript level of *CsF3’H-a*. Due to the low FPKM values (FPKM < 10) of *CsF3’H* and *CsF3’H-b*, these two transcripts were almost not expressed in fresh leave and solar-withered leaves, which was also confirmed by qRT-PCR. These results indicate that *CsF3’H-a* may function as the predominant transcript involved in flavonoid biosynthesis. The qRT-PCR analysis further confirmed the reliability of transcriptome datasets (Figure 6C). Earlier investigations have clearly demonstrated that the expression levels of flavonoid biosynthesis-related genes were closely related to the accumulation of flavonoids and catechins [44,45]. To identify these AS transcripts related to biosynthesis of flavonoids and catechins under different shading rates of solar-withering, the contents of flavonoids and catechins in all five samples were examined (Figure 6D). The contents of total flavonoids decreased sharply from FL to SW3, but exhibited a distinct rebound from SW3 to SW4. Likewise, solar-withering treatment continuously consumed the abundance of total catechins and eight individual catechin components, all of which showed the lowest value in SW3. Taken together, these results highlighted that reduction of flavonoids and catechins might result from the down-regulation of flavonoid biosynthesis-related genes and their AS transcripts.

Catechins, especially galloylated catechins, has positive effects on human health benefits and are the major contributors to bitter and astringent tastes [20,46]. Excessive catechins are detrimental to the formation of high-quality tea flavor. In turn, the drastic reduction of catechins attenuates the health benefits of tea. In this study, the next critical question was whether solar-withering process improved the palatability of oolong tea and diminished the health benefits of tea. Intriguingly, we noted that two genes, *ascorbate peroxidase1* (*APX1*) and *glutathione peroxidase3* (*GPX3*), involved in metabolic pathways underwent AS events. These two genes were reported to be responsible for the catalytic conversion of catechins into theaflavins [47,48]. Additionally, theaflavins are a group of high-molecular-weight polymeric compounds that endow tea with beneficial health-promoting properties and characteristic mellow taste [49]. Consequently, we suspected that solar-withering strongly inhibited catechin biosythesis, concomitant with an obvious elevation in the contents of theaflavins. To evaluate this possibility, the contents of total theaflavins and four individual theaflavin components obtained from metabolome were firstly analysis. As expected, substantial enhancements of all individual theaflavins and total theaflavins were observed under solar-withering treatment (Figure 6D). It was noted that the contents of theaflavins was inversely correlated with the that of catechins under solar-withering. Next, we found that the expression level of *CsGPX3*, *CsAPX1*, and *CsAPX1*-*a* were dramatically induced from FL to SW3, but showed a distinct decline at SW4. Distinguishingly, one AS transcript *CsGPX3*-*a* were insensitive to the environmental perturbation caused by solar-withering stage. We then observed a premature stop codon (PTC) in mRNA structure of *CsGPX3*-*a* (Figure 6E), implying that the occurrence of PTC may lead to the notably variations in structural and functional differences between full-length transcripts and their AS transcripts.

**Figure 6.**
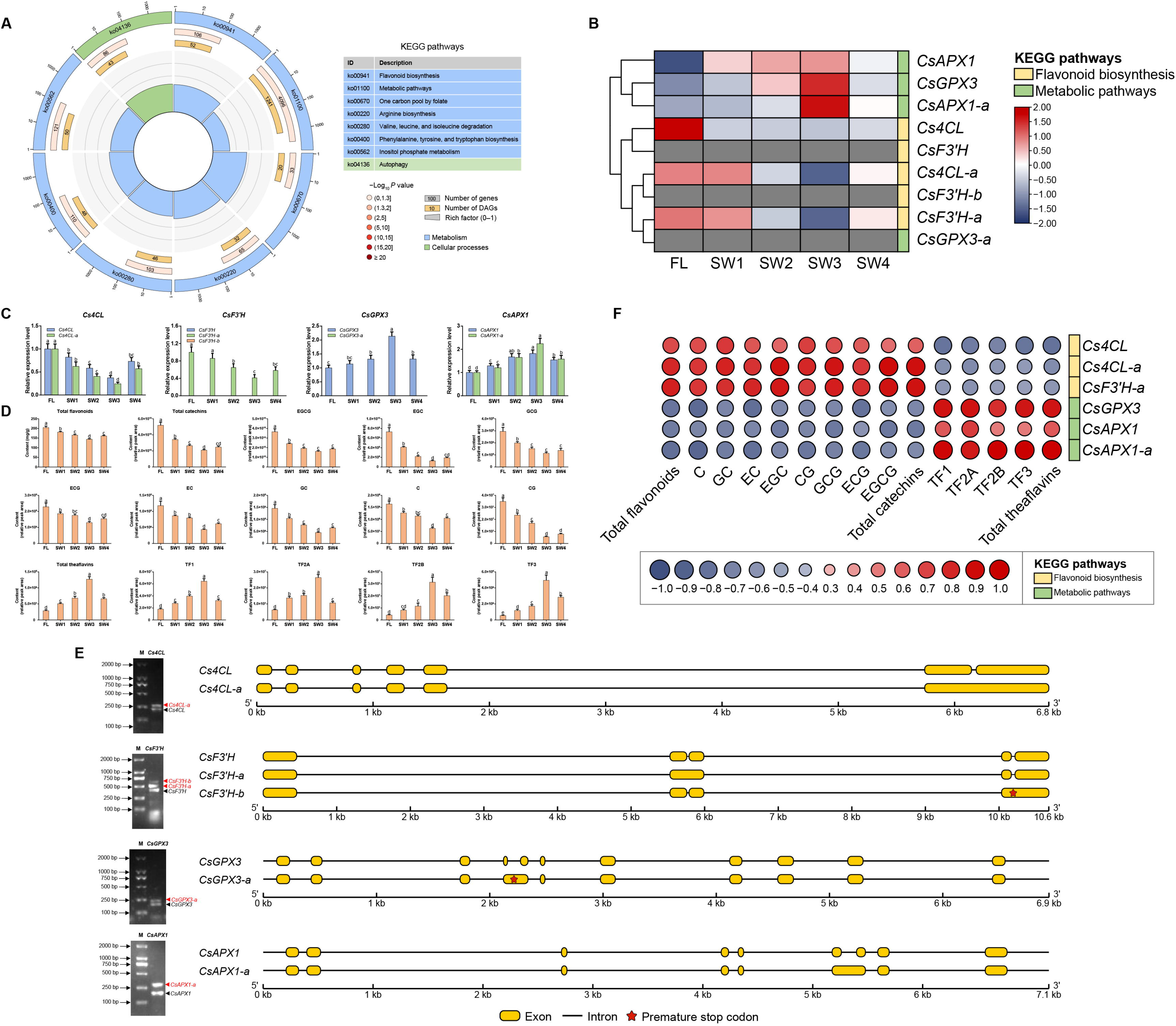
AS-mediated regulatory mechanism influences the accumulation of flavor-related metabolites. **A.** KEGG enrichment analysis of DAGs. From the outside to the inside, the first circle indicates enriched KEGG pathways and the number of the genes corresponds to the outer circle. The second circle indicates the number of the genes in the genome background and *P* value for enrichment of the genes for the specified KEGG pathway. The third circle indicates the number of the DAGs. The fourth circle indicates the enrichment factor of each KEGG pathway. **B.** The heatmap of DAGs and their AS transcripts under solar-withering with different shading rates based on the transcriptome datasets. The expression values of DAGs and their AS transcripts were normalized by log_2_-transformed (FPKM+1). Transcripts that are not expressed are shown in gray. **C.** Expression patterns of *Cs4CL*, *CsF3’H*, *CsAPX1*, *CsGPX3* and their AS transcripts under solar-withering with different shading rates determined by qRT-PCR. **D.** Relative contents of flavonoids, catechins, and theaflavins under solar-withering different shading rates determined by LC-MS. **E.** RT-PCR validation and gene structure of the full-length transcripts and AS transcripts. M indicates the lane with DNA size markers. The full-length transcripts and AS transcripts on the gel images are denoted with black and red triangles, respectively. **F.** Correlation analyses of full-length transcripts, AS transcripts, and flavor metabolites. The correlation analyses were performed based on Pearson’s correlation coefficient. A correlation coefficient greater than 0 indicates a positive correlation, while a correlation coefficient less than 0 indicates a negative correlation. Data are presented as mean ± standard deviation. Different letters indicate a significant difference (*P* < 0.05). FL, fresh leaves; SW1, solar-withering with high shading rate; SW2, solar-withering with middle shading rate; SW3, solar-withering with low shading rate; SW4, solar-withering with natural sunlight; AS, alternative splicing; DAGs, differentially expressed AS genes; KEGG, Kyoto encyclopedia of genes and genomes; FPKM, fragments per kilobase per million mapped reads; qRT-PCR, quantitative real-time polymerase chain reaction; LC-MS, liquid chromatography-mass spectrometry. RT-PCR, reverse transcription-polymerase chain reaction.

To better unveil the possible effects of AS transcripts on the accumulation of flavonoids, catechins, and theaflavins. the relationships between abundance of flavonoid-related AS transcripts and metabolites were lucubrated by Pearson’s correlation analysis (Figure 6F). Three transcripts, *CsF3’H*, *CsF3’H-b* and *CsGPX3*-*a*, were excluded from this analysis because they were not expressed in fresh leaves and solar-withered leaves. The expression levels of full-length transcript *Cs4CL* and its AS transcript *Cs4CL*-*a* were positively correlated with the contents of total flavonoids, total catechins, and eight individual catechins. Likewise, positive correlations were observed between the expression of AS transcript *CsF3’H*-*a* and the amount of above-mentioned metabolites. Moreover, we observed that the accumulation of total theaflavins and four individual theaflavin components showed positive correlations with *CsGPX3*, *CsAPX1*, and *CsAPX1*-*a*. We further examined the impacts of AS transcripts (*Cs4CL-a*, *CsF3’H-a*, and *CsAPX1-a*) on the accumulation of associated metabolites using an siRNA-mediated gene silencing strategy (Figure S10A). As expected, the transcript levels of *Cs4CL-a*, *CsF3’H-a*, and *CsAPX1-a* were considerably down-regulated in gene-silenced leaves, whereas the expression levels of these AS transcripts were not obviously altered by treatments with corresponding NC. Next, the flavonoid, catechin, and theaflavin in gene-silenced leaves were evaluated at the metabolic levels (Figure S10B). The abundance of total flavonoids, total catechins, and eight individual catechin components were vastly plummeted after *Cs4CL-a-* or *CsF3’H-a-*silenced treatment, while the amount of total theaflavins and four individual theaflavin components did not significantly change. Transcriptional repression of *CsAPX1-a* did not affect the contents of flavonoids and catechins in gene-silenced leaves, but dramatically hindered the accumulation of theaflavins. These data implied that some AS transcripts, instead of their corresponding full-length transcripts, are likely to play the predominant roles in the regulation of flavonoid, catechin, and theaflavin metabolisms. Additionally, the AS transcripts containing PTC may lose their original functions due to the remarkable variation in their mRNA structure. Overall, RNA methylation may lead to the dramatic alterations in flavonoid, catechin, and theaflavin metabolisms by triggering the AS regulatory mechanism, thus indirectly affecting the palatability of oolong tea.

## Discussion

### Dynamic changes in global m^6^A levels of tea leaves are mainly controlled by m^6^A erasers under solar-withering

In past decades, a series of studies on environmental stresses derived from tea processing, such as tea-withering stage, have greatly enriched our understanding of the molecular basis underlying specialized metabolisms in tea flavor formation [15,17]. Recently, it was elucidated that the tea flavor formation was implicated in the epigenetic modification, including DNA methylation and histone modification [28,29]. However, it remains unclear whether RNA methylation (epitranscriptome) participates in the regulation of flavor-related metabolisms and tea flavor formation during tea-processing. In the present study, we observed that m^6^A modifications are widely distributed on mRNAs in tea plant, and display dynamic changes in the m^6^A levels under solar-withering. After solar-withering treatment, the overall m^6^A levels were dramatically decrease in solar-withered leaves compared with fresh leaves. From FL to SW3, the overall m^6^A levels gradually decreased, but exhibited a distinct rebound from SW3 to SW4. These data indicated that solar-withering triggered a significant decrease in m^6^A levels, concomitant with a positive correlation between shading rate and globally m^6^A abundance within a certain range. According to earlier investigations [50,51], the overall DNA methylation level could be dynamically mediated by both DNA methyltransferases and demethylases, which catalyze the installation and excision of DNA methylation, respectively. Analogously, it was previously found that corresponding RNA methyltransferases and demethylases also exist in tea plant [14]. We therefore speculate that overall m^6^A abundance may be governed by RNA methyltransferases and demethylases under solar-withering. To evaluate this hypothesis, we comprehensive examined the expression profiles of these m^6^A regulatory genes under solar-withering. Notably, two m^6^A eraser genes, *CsALKBH4A* and *CsALKBH4B*, showed obvious changes at the transcript level under different shading rates of solar-withering, whereas the expression levels of other m^6^A eraser and reader members were altered slightly under solar-withering, consistent with the phenomenon in *Arabidopsis* [52] and tomato [11]. Similarly, during the solar-withering stage, tea leaves are subjected to multiple environmental stresses, including UV radiation [15]. Thus, our results implied that removal of m^6^A marks, rather than the installation and decoding of these marks, may play a dominant role under multiple stresses inflicted by tea-withering stage. During solar-withering with the shade treatment, excessive UV is filtered out and tea leaves are subjected to lower doses of UV radiation. In this case, the tea leaves are still alive and not seriously damaged. From FL to SW3, the transcription levels of *CsALKBH4A* and *CsALKBH4B* increased continuously. Furthermore, negative relationships between globally m^6^A abundance and the expression of two m^6^A erasers, as well as the increased overall m^6^A levels detected in *CsALKBH4*-silenced leaves further confirm that m^6^A demethylation is the major driving force for the decline in global m^6^A abundance from FL to SW3. However, tea leaves were exposed to high doses of UV radiation during solar-withering without the sunshade net (SW4). High doses of UV radiation are likely to have caused the plasma membrane to irreversibly lose its osmotic regulatory ability [53–55], leading to the cell rupture and further impeding the normal expression of nuclear-localized *CsALKBH4A* and *CsALKBH4B* [14]. Analogously, low doses of UV radiation induced *CsALKBH4* expression in a certain range during solar-withering with the sunshade net, while high doses of UV radiation strongly inhibited *CsALKBH4* expression during solar-withering without the sunshade net. The down-regulation of *CsALKBH4* expression in SW4 may impair the ability of m^6^A erasers to remove m^6^A modifications. This may be the reason why the overall m^6^A level exhibited an obvious rebound from SW3 to SW4. Collectively, dynamic changes in global m^6^A levels of tea leaves are mainly controlled by m^6^A erasers under different shading rates of solar-withering, and CsALKBH4-mediated RNA demethylation are more likely to play critical roles in the tea-withering stage.

### RNA demethylation directly contributes to the accumulation of volatile terpenoids and the formation of tea aroma

It is well-recognized that environmental stresses can greatly regulate the accumulation of specialized metabolites in tea leaves, thereby affecting tea quality [56]. During postharvest processing, the tea leaves inevitably encounter various environmental stresses, concomitant with obvious alterations in a large number of flavor-related compounds, which endows tea with unique flavor [17]. Although many efforts have been made to explore the effects of specialized metabolites on tea flavor formation at the transcriptional, translational and metabolic levels, the functional roles of epigenetic modification, especially RNA methylation, and precise regulatory mechanisms underlying the m^6^A-mediated flavor formation in the tea-withering stage remain vague.

In the current study, integrated RNA methylome and transcriptome analyses reveal that the variations in the number of DMPs under solar-withering with different shading rates were in line with the dynamics of total m^6^A abundance. The amount of DMPs continued to decreased from FL *vs.* SW1 to FL *vs.* SW3 comparisons, whereas the number of DMPs increased in the FL *vs.* SW4 comparison. The increase in the number of DMPs in FL *vs*. SW4 may be associated with the inhibition of *CsALKBH4* expression by high doses of UV radiation under no-shading treatment. The down-regulation of *CsALKBH4* expression in SW4 hindered RNA demethylation, thereby amplifying differences in m^6^A abundance between SW4 and other samples, and finally leading to an increase in the number of DMPs. Next, a total of 1137 genes containing DMPs under solar-withering treatments. KEGG pathway analysis showed that the identified DMP-associated genes were mainly enriched in terpenoid biosynthesis pathway. It is well documented that volatile terpenoids are major quality components of oolong tea, and have significant influences on the formation of floral and honey aroma in high-quality oolong tea due to their low odor thresholds [15]. Moreover, several lines of evidence suggest that environmental stresses can alter m^6^A levels of mRNA transcripts [12,57]. In our research, the expression levels of seven terpenoid biosynthesis-related genes were significantly induced under solar-withering, concomitant with remarkably elevations in volatile terpenoids. The positive correlations between expression levels of these structural genes related to terpenoid biosynthesis and terpenoid abundance implied that multiple stresses induced by solar-withering promotes the terpenoid accumulation by up-regulating terpenoid biosynthesis-related genes, consistent with the results of another study [18]. In addition, we discovered that m^6^A abundance of these terpenoid biosynthesis-related genes was negatively correlated with their corresponding expression levels, analogously to a previous report that genes with low m^6^A abundance tend to show up-regulated expression [58]. These results accordingly raise an open question that the expression levels of m^6^A-containing genes may be governed by RNA methylation. According to recent reports, knockout of m^6^A writers dramatically decreased m^6^A levels of m^6^A-modified genes, but significantly increased their expression levels [6,59]. Nevertheless, overaccumulation of m^6^A marks on mRNAs reduced their expression in the *AtALKBH10B* mutant line, suggesting that m^6^A modification had a negative effect on mRNA expression [60]. Consistently, the suppression of *CsALKBH4A* and *CsALKBH4B* resulted in a significant decrease in mRNA stability of *CsDXS* and three DMP-associated genes, concomitant with an obvious decline in their mRNA abundance. Simultaneously, the eight volatile terpenoids displayed a noticeable decrease in the *CsALKBH4A*- or *CsALKBH4B*-silenced leaves. These results support the speculation that, combined with the CsALKBH4-mediated RNA demethylation mentioned above, RNA demethylation may not only mediate overall m^6^A level by regulating the expression of m^6^A erasers but also enhance the mRNA stability of DMP-associated genes, thereby increasing their expression. However, some previous studies have found a positive correlation between the mRNA abundance and the intensity of m^6^A modification [13,61], which is contrary to our findings. Therefore, the mechanism underlying the regulatory effect of m^6^A modification on mRNAs under solar-withering stage needs in-depth dissection. A recent study unveiled that abiotic stress leads to the relocation of m^6^A enrichment on mRNA transcripts [9]. Data from this study showed that m^6^A peaks in *CsDXS* and other three DMP-associated genes were all distributed within 3′UTR and around stop codon under solar-withering, indicating that solar-withering only affects m^6^A levels of these DMP-associated genes but did not redistribute m^6^A marks on these genes. Revisiting the reports that m^6^A modification has a positive effect on mRNA abundance, we noted that these m^6^A marks are mainly concentrated in the CDS region. Combining the observation in our research, it is reasonable to assume that m^6^A modification may portray distinct regulatory roles in mRNA abundance based on the distribution of m^6^A modifications in mRNA structure. Taken together, CsALKBH4-mediated RNA demethylation promoted the expression of DMP-associated genes involved in terpenoid biosynthesis and accumulation of aroma-related terpenoids by enhancing their mRNA stability. Additionally, it is worth noting that shading rate was negatively correlated with the expression of terpenoid biosynthesis-related genes and terpenoid contents from FL to SW3, implying that, within a certain range, the moderate shading treatment helped promote terpenoid biosynthesis and the formation of a high-quality aroma in oolong tea. Inadequate shading rate in SW4 arrested CsALKBH4-mediated RNA demethylation, thereby destabilizing the terpenoid biosynthesis-related genes, and ultimately leading to a significant decrease in the terpenoid abundance. This was also in line with the actual production, moderate shading treatment was beneficial to improve the tea flavor. These findings indicate the feasibility of improving solar-withering through controlled the shading rate of solar-withering, and the shading threshold is a critical indicator to improve tea aroma.

### RNA modification indirectly affects the contents of flavonoids, catechins and theaflavins by triggering the AS regulatory mechanism

AS regulatory mechanism is well-known as a crucial strategy for plants to respond to diverse environmental stresses [62]. By generating a large number of AS transcripts, the diversity of coding proteins can be enhanced to alleviate the adverse effects of stresses. Although many AS events were detected in the secondary metabolism, the specific regulatory mechanism of AS events on flavor-related genes and the formation of tea flavor quality during tea processing remains elusive. Furthermore, current studies illustrate that m^6^A modification has important regulatory effects on RNA splicing [40]. Likewise, we observed that CsALKBH4-mediated RNA demethylation contributed to *CsSm* abundance in the spliceosome pathway via enhancement of its mRNA stability. According to previous research, *Sm* is regarded as a crucial component of the spliceosome [63]. Its main function is to assemble onto several small nuclear RNAs (snRNAs), and then bind with a series of additional proteins to form small nuclear ribonucleoprotein (snRNP) particles. Within the whole spliceosome, the snRNA makes a decisive contribution to catalysis and recognition of splicing sites in pre-mRNA splicing. Intriguingly, both the frequency of AS events and the number of DAGs were closely associated with the *CsSm* expression. We therefore inferred that RNA demethylation might be involved in the regulation of AS events by modulating the m^6^A abundance and expression levels of spliceosome-related genes.

To further investigate the functional roles of AS-mediated regulation under solar-withering, we conducted KEGG annotation on all identified DAGs. The majority of DAGs were enriched in metabolic pathways and flavonoid biosynthesis pathway. Further expression analysis of four DAGs involved in these two pathways revealed that the AS transcripts encompassing PTC (*CsF3’H*-*b* and *CsGPX3*-*a*) were almost not expressed under solar-withering. This phenomenon may be due to the introduction of PTC into the gene structure, leading in the production of loss-of-function truncated proteins, and these truncated proteins may then be degraded via the nonsense-mediated decay (NMD) pathway. Furthermore, this speculation is supported by correlation analysis between metabolite accumulation and the expression of AS transcripts, which showed that *CsF3’H*-*b* were not correlated with the accumulation of flavonoid and catechin. Also, *CsGPX3*-*a* were marginally correlated with the theaflavin level. It is noted that the expression patterns of two non-PTC type AS transcripts, *Cs4CL-a* and *CsF3’H-a,* were consistent with those of relevant full-length transcript under solar-withering, respectively. Meanwhile, *Cs4CL-a* and *CsF3’H-a* showed higher expression levels relative to their full-length transcripts at each shading rate of solar-withering. These findings implied that these two AS transcripts act as the predominant transcripts in the flavonoid biosynthesis pathway. As a precedent for this hypothesis, the *CsbHLH-2*, an AS transcript, is predominately formed under cold stress and enhances stress tolerance through positive regulation of some signaling pathways [64]. Another AS transcript *CsAPX1*-*a* may coordinate with its full-length transcript to regulate theaflavin biosynthesis. Aforementioned speculation was further strengthened by our correlation analysis of AS transcripts and metabolites under solar-withering, which revealed that *Cs4CL-a* and *CsF3’H-a* were positively correlated the accumulation of flavonoids and catechins. The positive correlations were also detected between AS transcript *CsAPX1*-*a* and theaflavin content. We inferred that the accumulation of flavonoids, catechins, and theaflavin was mediated by not only canonical full-length transcripts but also extraordinary AS transcripts, which may serve as major coordinators at the post-transcriptional level under solar-withering. Therefore, the down-regulation of *Cs4CL*, *Cs4CL*-*a*, and *CsF3’H*-*a* as well as up-regulation of *CsGPX3*, *CsAPX1*, and *CsAPX1-a* in solar-withered leaves synergistically dampened the flavonoid biuosynthesis and expedited the flavonoid catabolism, jointly promoting the conversion of bitter and astringent substances (flavonoid and catechin) into mellow-tasting theaflavin. Compared with other shading rates of solar-withering, SW3 had the lowest content of catechins and high content of theaflavins, indicating that moderate shading rate, rather than insufficient or excessive shading rate, could effectively reduce bitterness and astringency and enhance the mellow taste of tea infusions. Together, these findings highlighted that RNA demethylation indirectly affect the accumulation of flavonoids, catechins and theaflavins by triggering the AS-mediated regulatory mechanism, thereby improving the palatability of oolong tea.

In conclusion, integrated RNA methylome and transcriptome analyses reveal that m^6^A-mediated regulatory mechanism act as the central coordinators in the accumulation of tea specialized metabolites under solar-withering. These results showed that dynamic changes in global m^6^A levels of tea leaves are mainly controlled by m^6^A erasers under different shading rates of solar-withering. Moreover, CsALKBH4-driven RNA demethylation can not only directly affect the accumulation of volatile terpenoids and the tea aroma formation by mediating the stability and abundance of terpenoid biosynthesis-related genes, but also indirectly regulate the contents of flavonoids, catechins, and theaflavins, as well as the tea taste formation via triggering the AS-mediated regulatory mechanism. These findings uncovered a novel layer of epitranscriptomic gene regulation in tea flavor-related metabolic pathways and established a strong link between m^6^A-mediated regulatory mechanism and the improvement of high-quality flavor and palatability in oolong tea (**Figure 7**). Our work not only provided a solid foundation for deciphering the functional roles of m^6^A modification in tea plant, but also broadened our understanding of regulatory mechanisms underlying the m^6^A-mediated flavor formation under solar-withering.

**Figure 7.**
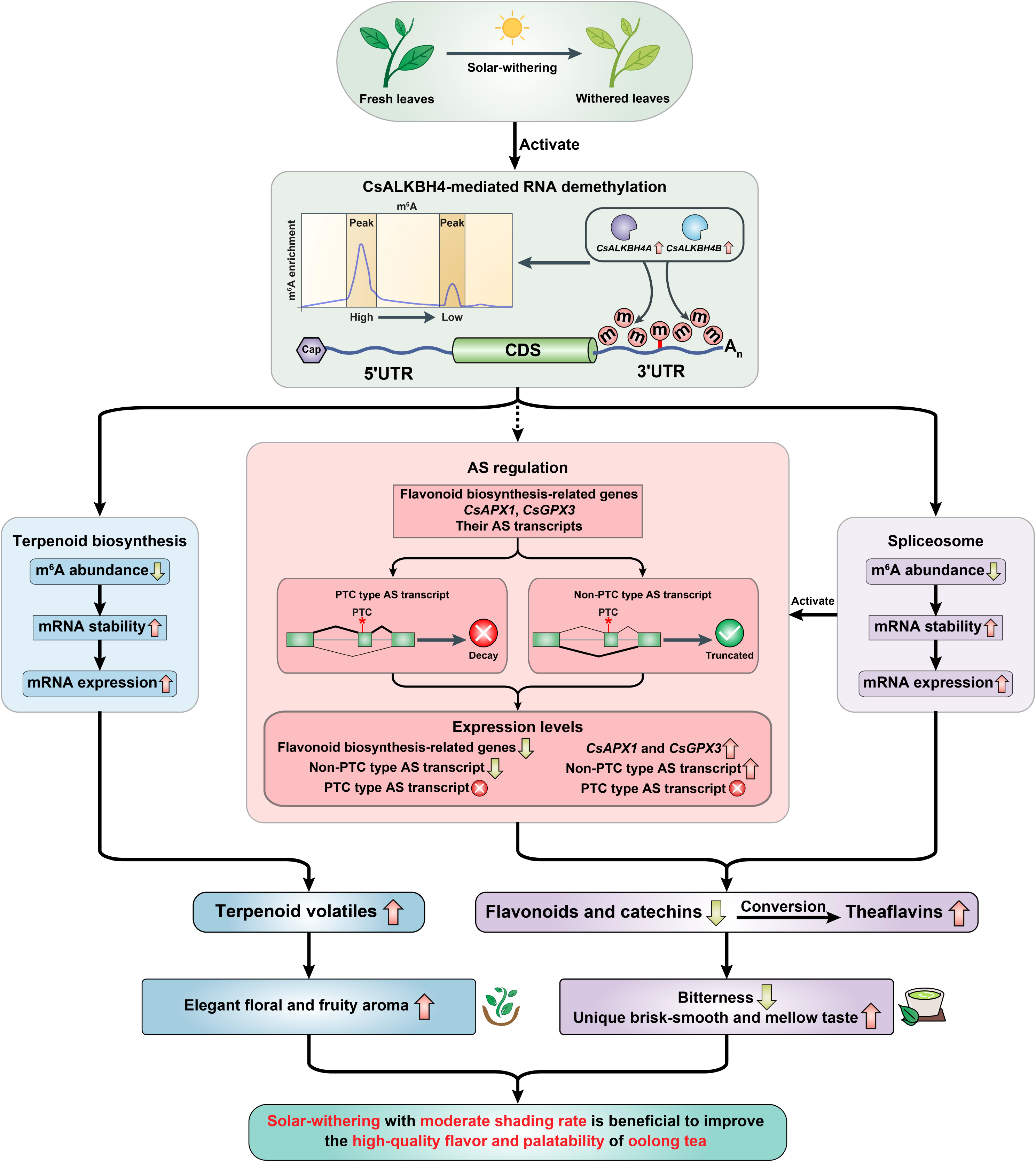
Schematic model for the effects of m^6^A-mediated regulatory mechanism on the accumulation of flavor metabolites in tea (*Camellia sinensis*) leaves under solar-withering. CsALKBH4-driven RNA demethylation can not only directly affect the accumulation of volatile terpenoids and the tea aroma formation by mediating the stability and abundance of terpenoid biosynthesis-related genes, but also indirectly regulate the contents of flavonoids, catechins, and theaflavins, as well as the tea taste formation via triggering the AS-mediated regulatory mechanism. These findings uncovered a novel layer of epitranscriptomic gene regulation in tea flavor-related metabolic pathways and established a strong link between m^6^A-mediated regulatory mechanism and the improvement of high-quality flavor and palatability in oolong tea. Solid arrows indicate direct regulation, and dashed arrows indicate indirect regulation. 5′UTR, 5′ untranslated region; CDS, coding sequence; 3′UTR, 3′ untranslated region; AS, alternative splicing; PTC, premature stop codon.

## Materials and methods

### Plant materials and solar-withering treatments

Fresh tea bud with first three leaves (*Camellia sinensis* cv. Tieguanyin) were obtained from a tea plantation located at Fujian Agriculture and Forestry University, Fuzhou, China (E 119°14′, N 26°05′). The picked tea leaves were evenly divided into five portions. The first portion of fresh leaves was sampled immediately without any processing. The other four portions were laid out on bamboo sieves. For solar-withering treatments with different shading rates, black nylon sunshade nets with different meshes were placed 0.2 m over the bamboo sieves. The detailed withering treatments were performed as follows: solar-withering with high shading rate (SW1, sunshade net with 1500 meshes, 10,000 ± 400 lx); solar-withering with middle shading rate (SW2, sunshade net with 1000 meshes, 20,000 ± 800 lx); solar-withering with low shading rate (SW3, sunshade net with 500 meshes, 40,000 ± 1200 lx); solar-withering with natural sunlight (SW4, without sunshade net, 80,000 ± 2000 lx). The duration of solar-withering was 45 min, and other environmental parameters were consistent with our previous report [24]. Three independent biological replicates were performed for each solar-withering treatment. The light intensity and spectrum were measured by the spectral irradiance colorimeter (Catalog No. SPIC-300, Everfine Corporation, Hangzhou, China). Simultaneously, UV intensity was recorded by the UV radiometer (Catalog No. UV340B, Sanpometer Corporation, Shenzhen, China). After solar-withering, all the samples were collected and frozen immediately in liquid nitrogen. Then, the frozen samples were maintained at −80 °C until further investigation.

### Quantitative analysis of m^6^A/A ratio

Global m^6^A/A ratio in tea leaves was determined as previously reported [32]. In brief, RNA was extracted from each sample using the TransZol UP Reagent (Catalog No. ET111, TransGen, Beijing, China). Then, eligible RNA was used to conducted quantitative analysis of global m^6^A abundance by EpiQuik m^6^A RNA Methylation Quantification Kit (Catalog No. P-9005, Epigentek, Farmingdale, NY), following the manufacturer’s instructions.

### MeRIP-seq and data analysis

For methylated RNA immunoprecipitation sequencing (MeRIP-seq), total RNA was extracted separately from each sample using the TransZol UP Reagent (Catalog No. ET111, TransGen). The integrity and quantity of obtained RNA were assessed by RNase-free agarose gel electrophoresis and an Agilent 2100 Bioanalyzer platform (Agilent Technologies). All RNA samples passed quality checks, and no signs of RNA degradation were found in fresh leaves and four solar-withered leaves with different shading rates (Figure S11). Next, poly (A) mRNA was isolated from qualified RNA using the Dynabeads mRNA Purification Kit (Catalog No. 61,006, Thermo Fisher Scientific). Then, the enrichment of m^6^A were performed using the Magna MeRIP m^6^A Kit (Catalog No. 17-10,499, Millipore, Billerica, MA). In brief, the mRNA was fragmented into about 100 nucleotide-long fragments by using fragmentation buffer. The fragmented RNA was divided into two portions, one of which was enriched with m^6^A-specific antibody in IP buffer. The other RNA without IP treatment was used as the input control. The m^6^A-containing fragments and non-immunoprecipitated fragments were used for strand-specific library construction with NEBNext Ultra II RNA Library Prep Kit (Catalog No. E7770, NEB, Ipswich, MA). The average insertion size of each paired-end library was 100 ± 50 bp. Finally, m^6^A MeRIP sequencing were performed using an Illumina NovaSeq 6000 platform (Illumina, San Diego, CA). Three independent biological replicates for each sample were sequenced.

The raw reads containing adapter sequences and undetermined bases were trimmed out by Trimmomatic tool [65]. After trimming, high-quality reads were mapped to the chromosome-level genome of tea plant using HISAT software [66]. R-package exomePeak [67] was used to identify the m^6^A peak in each m^6^A-immunoprecipitated sample with the corresponding non-immunoprecipitated sample acting as a background. In all three independent biological samples, peaks with overlap more than 50% of their length and *P* < 0.05 were designated as high-confidence m^6^A peaks. The visualization of m^6^A peaks were was performed using Integrative Genomics Viewer software [68]. According to the genomic location information, the distribution of peak on non-overlapping mRNA regions, such as 5′UTR, CDS and 3′UTR, was determined using BedTools [69]. The MEME suite [70] and HOMER tool [71] were used for motif search within m^6^A peaks. DMPs between groups were identified using DiffBind software [72] with |FC| ≥ 2 and *P* < 0.05 as the thresholds. The expression of mRNAs from input libraries was calculated as the FPKM value by using StringTie software [73]. DEGs were identified according to the criteria of |FC| ≥ 2 and *P* < 0.05 by using DESeq2 software [74]. The expression profiles of candidate genes were visualized using the TBtools software [75] based on the standardized FPKM values. The rMATS software was employed to detect the AS events. AS events with false discovery rate (FDR) < 0.05 were considered significant AS events. The AS transcripts were validated as described previously [76] using reverse transcription-polymerase chain reaction (RT-PCR) with specific primers (Table S11). The PCR products were then monitored by agarose gel electrophoresis. DEGs that underwent AS events were defined as DAGs according to previous criteria [64]. KEGG enrichment analysis of DMP-associated genes and DAGs was performed as previously reported [25].

### m^6^A-IP-qPCR and qRT-PCR

The m^6^A-IP-qPCR was referred to previous method [11] with minor modifications. In brief, the RNAs for m^6^A MeRIP-seq was fragmented into about 300 nucleotide-long fragments by using the aforementioned fragmentation buffer. Part of the fragmented RNAs was used for m^6^A-IP using m^6^A specific antibody. Another part of RNA without IP treatment was used as the input control. The m^6^A-containing RNAs and non-immunoprecipitated RNAs were reversely transcribed into cDNAs using the TransScript First-Strand cDNA Synthesis SuperMix Kit (Catalog No. AT301, TransGen). The m^6^A enrichment in specific mRNA region was detected by using the LightCycler 480 platform (Roche, Basel, Switzerland). The abundance of m^6^A enrichment was quantified by using the 2^−ΔΔCT^ method [77]. Relative abundance of specific mRNA region for the m^6^A-IP sample was firstly standardized against that of *CsActin* (GenBank accession: HQ420251), which has no obvious m^6^A-modified peak and was served as an internal control, and then standardized against that for the non-immunoprecipitated sample.

The qRT-PCR was performed on the LightCycler 480 platform (Roche) according to our previously described method [24]. The RNAs without fragmentation treatment were reversely transcribed into cDNAs as described above. *CsActin* (GenBank accession: HQ420251) was used to normalize the mRNA expression. Relative mRNA levels were calculated using 2^−ΔΔCT^ method. Three independent biological replicates were performed for each assay. All specific primers used for m^6^A-IP-qPCR and qRT-PCR were shown in Table S11.

### Volatile analysis by GC-MS

The extraction and analysis of volatile compounds in tea samples were performed as previously described [78]. An Agilent Model 7890B chromatograph-mass spectrometer (Agilent Technologies, Palo Alto, CA) equipped with a 7000D mass spectrometer (Agilent Technologies) were utilized to detect the volatile compounds. Three replicates of each assay were conducted. The detected volatiles were identified according to the retention times and mass spectra data in National Institute of Standards and Technology Mass Spectral Library. Relative abundance of individual volatile compound was shown by chromatographic peak area.

### Metabolomic analysis by LC-MS

The metabolite extraction and metabolomic analysis were essentially as described previously [79]. Briefly, the freeze-dried tea samples were ground into powder using a tissue grinder (Catalog No. JXFSTPRP-24, Jingxin Company, Shanghai, China). Then, the powdered samples were extracted with 800 μL of 70% methanol and 250 µL 2′,7′-dichlorofluorescein, followed by a centrifugation at 14,000 *g* for 15 min. The supernatant was passed through a 0.22-µm polyvinylidene fluoride filter and analyzed using the Acquity two-dimensional ultra-performance liquid chromatography (2D-UPLC) platform (Waters, Milford, MA) connected to a Q-Exactive quadrupole-orbitrap mass spectrometry (Thermo Fisher Scientific, Waltham, CA). Separation of compounds was achieved on a Hypersil GOLD aQ column (100 mm × 2.1 mm, 1.9 μm, Thermo Fisher Scientific) with 0.1% formic acid in pure water (v/v, solvent A) and 0.1% formic acid in acetonitrile (v/v, solvent B) at a flow rate of 0.3 mL/min. The gradient elution was initiated with 5% solvent B for 2 min, linearly elevated to 95% solvent B over 22 min, kept at 95% solvent B for 5 min, then return to the initial conditions (5% solvent B) within 3 min. The column temperature was set to 40 °C. Peak detection and retention time correction were performed by the software Compound Discoverer 3.1 (Thermo Fisher Scientific). The detected metabolites were identified by comparing their molecular mass, retention time, and mass spectrometry fragmentation patterns with those of the authentic standards and standard databases (mzCloud and mzVault). Relative abundance of each metabolite was calculated using metaX tool [80]. Moreover, the total flavonoid contents of tea leaves were detected according to aluminum chloride colorimetric method [24]. Three independent biological replicates were performed for each experiment.

### Gene suppression and mRNA stability assays

Gene suppression assay was performed as previously described [14]. The freshly detached tea bud and first leaf from natural grown tea plants were incubated in 1.5 mL microcentrifuge tubes that contained 1 mL of 20 μM siRNA or siRNA-negative control (siRNA-NC) solution. After incubation for 12 h and 24 h, the tea bud and first leaf were harvested and then used for qRT-PCR. The specific siRNAs were obtained from GenePharma (Shanghai, China). Detail information of siRNA and siRNAs-NC is listed in Table S11. Overall m^6^A abundance and metabolite content of gene-silenced leaves and NC-leaves were examined as described above.

To assess the mRNA stability, the leaf discs were collected from gene-silenced tea leaves and incubated in sterile water that contained 10 μg/mL actinomycin D (Catalog No. A1410, Sigma) solution. Tea leaves incubated in sterile water were as controls. Total RNAs were isolated from the leaves sampled at 6 h and 12 h, respectively. The mRNA stability was determined according to a reported method [32] by qRT-PCR using specific primers (Table S11).

### Statistical analyses

The correlation analyses of full-length transcripts, AS transcripts and flavor metabolites were performed based on Pearson’s correlation coefficient. The difference among various groups were conducted by using one-way analysis of variance followed by Tukey’s *post hoc* test. Data were shown as the mean ± standard deviation. The *P* < 0.05 was considered statistically significant. The raw data for qRT-PCR are supplied as Table S12.

## Data availability

The raw sequencing data have been deposited to the Sequence Read Archive (SRA) at the National Center for Biotechnology Information (NCBI) under the accession number PRJNA762445 and Genome Sequence Archive (GSA) at the National Genomics Data Center (NGDC) under the accession number CRA006400.

## CRediT author statement

**Chen Zhu:** Methodology, Validation, Formal analysis, Investigation, Writing - original draft, Writing - Review & Editing. **Shuting Zhang:** Software, Formal analysis, Data curation, Writing - original draft. **Chengzhe Zhou:** Validation, Investigation. **Caiyun Tian:** Validation, Investigation. **Biying Shi:** Software, Data curation. **Kai Xu:** Validation, Investigation. **Linjie Huang:** Validation, Investigation. **Yun Sun**: Software, Data curation. **Yuling Lin:** Conceptualization, Formal analysis. **Zhongxiong Lai:** Conceptualization, Formal analysis, Writing - review & editing, Writing - Review & Editing, Funding acquisition, Project administration. **Yuqiong Guo:** Conceptualization, Methodology, Formal analysis, Investigation, Writing - review & editing, Writing - Review & Editing, Project administration.

## Competing interests

The authors have declared no competing interests.

## Acknowledgments

This work was supported by the Earmarked Fund for China Agriculture Research System of MOF and MARA (Grant No. CARS-19), the Scientific Research Foundation of Graduate School of Fujian Agriculture and Forestry University (Grant No. 324-1122yb070), the Scientific Research Foundation of Horticulture College of Fujian Agriculture and Forestry University (Grant No. 2019B01), the Rural Revitalization Tea Industry Technical Service Project of Fujian Agriculture and Forestry University (Grant No. 11899170145), The “Double first-class” scientific and technological innovation capacity and enhancement cultivation plan of Fujian Agriculture and Forestry University (Grant No. KSYLP004), 6.18 Tea Industry Technology Branch of Collaborative Innovation Institute (Grant No. K1520001A), Fujian Agriculture and Forestry University Construction Project for Technological Innovation and Service System of Tea Industry Chain (Grant No. K1520005A01), the Construction of Plateau Discipline of Fujian Province (Grant No. 102/71201801101), Tea Industry Branch of Collaborative Innovation Institute of Fujian Agriculture and Forestry University (Grant No. K1521015A), and the Special Fund for Science and Technology Innovation of Fujian Zhang Tianfu Tea Development Foundation (Grant No. FJZTF01). We are grateful to Timur Zaripov from Fujian Agriculture and Forestry University for his linguistic assistance.

## Supporting material

**Figure S1 The light intensity, spectrum, and UV intensity of solar-withering with different shading rates**

SW1, solar-withering with high shading rate; SW2, solar-withering with middle shading rate; SW3, solar-withering with low shading rate; SW4, solar-withering with natural sunlight; UV, ultraviolet.

**Figure S2 Venn diagrams showing the overlap of m^6^A peaks identified from fresh leaves and four solar-withered leaves with different shading rates**

FL, fresh leaves; SW1, solar-withering with high shading rate; SW2, solar-withering with middle shading rate; SW3, solar-withering with low shading rate; SW4, solar-withering with natural sunlight

**Figure S3 Venn diagrams showing the overlap of m^6^A-marked genes identified from fresh leaves and four solar-withered leaves with different shading rates**

FL, fresh leaves; SW1, solar-withering with high shading rate; SW2, solar-withering with middle shading rate; SW3, solar-withering with low shading rate; SW4, solar-withering with natural sunlight

**Figure S4 Enrichment score of m^6^A peaks in five transcript segments**

FL, fresh leaves; SW1, solar-withering with high shading rate; SW2, solar-withering with middle shading rate; SW3, solar-withering with low shading rate; SW4, solar-withering with natural sunlight; 5′UTR, 5′ untranslated region; CDS, coding sequence; 3′UTR, 3′ untranslated region.

**Figure S5 Correlation between m6A modification and expression levels of the DMP-associated genes under solar-withering with different shading rates**

FL, fresh leaves; SW1, solar-withering with high shading rate; SW2, solar-withering with middle shading rate; SW3, solar-withering with low shading rate; SW4, solar-withering with natural sunlight; DMP-associated genes, differentially methylated peak-associated genes.

**Figure S6 m^6^A abundance and expression levels of DMP-associated genes under solar-withering with different shading rates based on RNA methylome and transcriptome datasets**

**A.** m^6^A abundance of DMP-associated genes under solar-withering with different shading rates. **B.** Expression levels of DMP-associated genes under solar-withering with different shading rates. The m^6^A abundance and expression levels of DMP-associated genes were normalized by log_2_-transformed (m^6^A enrichment+1) and log_2_-transformed (FPKM+1), respectively. FL, fresh leaves; SW1, solar-withering with high shading rate; SW2, solar-withering with middle shading rate; SW3, solar-withering with low shading rate; SW4, solar-withering with natural sunlight; DMP-associated genes, differentially methylated peak-associated genes; FPKM, fragments per kilobase per million mapped reads.

**Figure S7 m^6^A abundance and expression levels of *CsRB-L10e*, *CsEIF4A*, *CsSm*, and *CsDXS* under solar-withering with different shading rates based on RNA methylome and transcriptome datasets**

**A.** m^6^A abundance of *CsRB-L10e*, *CsEIF4A*, *CsSm*, and *CsDXS* under solar-withering with different shading rates. **B.** Expression levels of *CsRB-L10e*, *CsEIF4A*, *CsSm*, and *CsDXS* under solar-withering with different shading rates. Data are presented as mean ± standard deviation. Different letters indicate a significant difference (*P* < 0.05). FL, fresh leaves; SW1, solar-withering with high shading rate; SW2, solar-withering with middle shading rate; SW3, solar-withering with low shading rate; SW4, solar-withering with natural sunlight; FPKM, fragments per kilobase per million mapped reads.

**Figure S8 Relative m^6^A enrichment and relative expression levels of *CsRB-L10e*, *CsEIF4A*, *CsSm*, and *CsDXS* in *CsALKBH4A*- or *CsALKBH4B*-silenced leaves**

**A.** Relative m^6^A enrichment of *CsRB-L10e*, *CsEIF4A*, *CsSm*, and *CsDXS* in *CsALKBH4A*- or *CsALKBH4B*-silenced leaves determined by m^6^A-IP-qPCR. **B.** Relative expression levels of *CsRB-L10e*, *CsEIF4A*, *CsSm*, and *CsDXS* in *CsALKBH4A*- or *CsALKBH4B*-silenced leaves determined by qRT-PCR. m^6^A-IP-qPCR, m^6^A-immunoprecipitation-qPCR; qRT-PCR, quantitative real-time polymerase chain reaction.

**Figure S9 Expression levels of DAGs and their AS transcripts involved in flavonoid biosynthesis pathway under solar-withering with different shading rates based on transcriptome datasets**

Expression values of DAGs and their AS transcripts were normalized by log_2_-transformed (FPKM+1). FL, fresh leaves; SW1, solar-withering with high shading rate; SW2, solar-withering with middle shading rate; SW3, solar-withering with low shading rate; SW4, solar-withering with natural sunlight; DAGs, differentially expressed AS genes; AS, alternative splicing; FPKM, fragments per kilobase per million mapped reads.

**Figure S10 Relative expression levels of *Cs4CL-a*, *CsF3’H-a*, and *CsAPX1-a*, as well as the contents of flavor metabolites in AS transcript-silenced leaves**

**A.** Relative expression levels of *Cs4CL-a*, *CsF3’H-a*, and *CsAPX1-a* in AS transcript-silenced leaves determined by qRT-PCR. **B.** The contents of flavor metabolites in AS transcript-silenced leaves determined by LC-MS. Data are presented as mean ± standard deviation. Single asterisk indicates a significant difference (*P* < 0.05). Double asterisks indicate a highly significant difference (*P* < 0.01). AS, alternative splicing; qRT-PCR, quantitative real-time polymerase chain reaction; LC-MS, liquid chromatography-mass spectrometry.

**Figure S11 RNA quality of fresh leaves and four solar-withered leaves with different shading rates**

Three independent biological replicates were performed for each sample. FL, fresh leaves; SW1, solar-withering with high shading rate; SW2, solar-withering with middle shading rate; SW3, solar-withering with low shading rate; SW4, solar-withering with natural sunlight.

**Table S1 A summary of m^6^A-seq reads in tea leaves under solar-withering with different shading rates**

**Table S2 The number of m^6^A peaks detected in m^6^A-marked genes**

**Table S3 Statistics of DMPs under solar-withering with different shading rates**

**Table S4 DMP-associated genes involved in ribosome pathway**

**Table S5 DMP-associated genes involved in RNA transport pathway**

**Table S6 DMP-associated genes involved in spliceosome pathway**

**Table S7 DMP-associated genes involved in terpenoid biosynthesis pathway**

**Table S8 Statistics of AS events under solar-withering with different shading rates**

**Table S9 Statistics of DAGs under solar-withering with different shading rates**

**Table S10 DAGs involved in flavonoid biosynthesis pathway**

**Table S11 Primers used in this study**

**Table S12 Raw data for qRT-PCR**

