## Supplementary figures and images for "RNA Methylome Reveals the m^6^A-Mediated Regulation of Flavor Metabolites in Tea Leaves under Solar-Withering"

### Figure S1

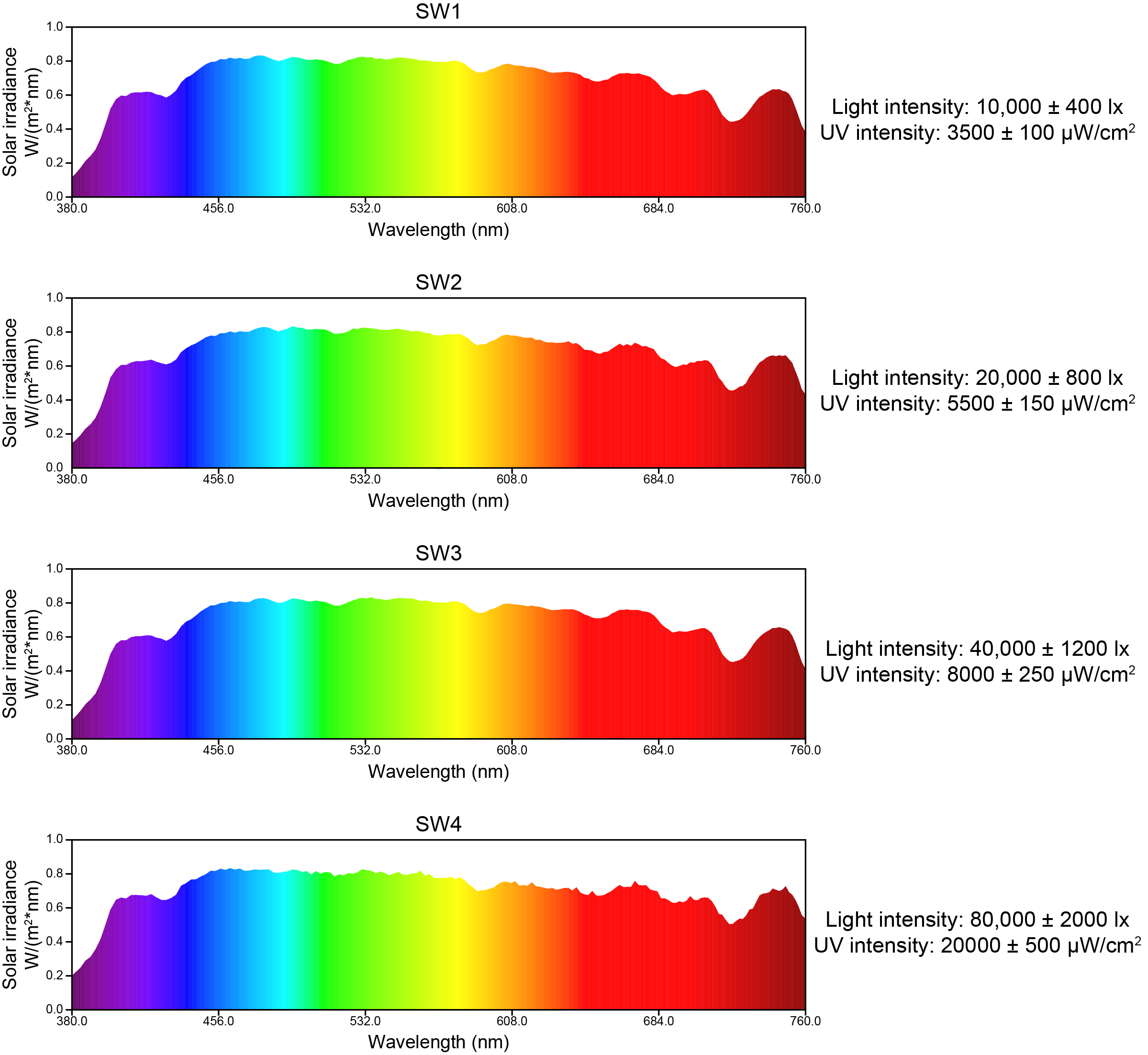

### Figure S2

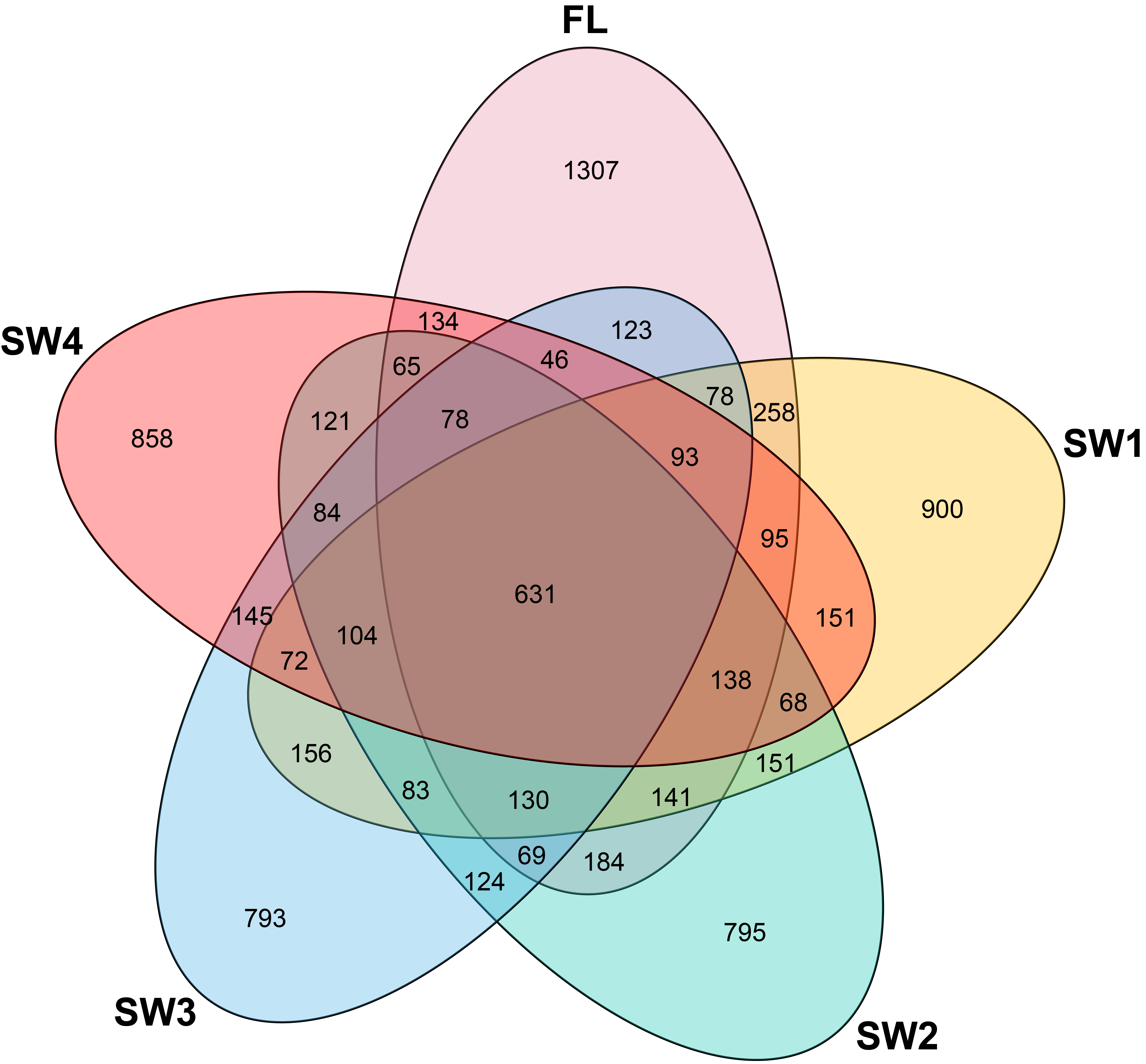

### Figure S3

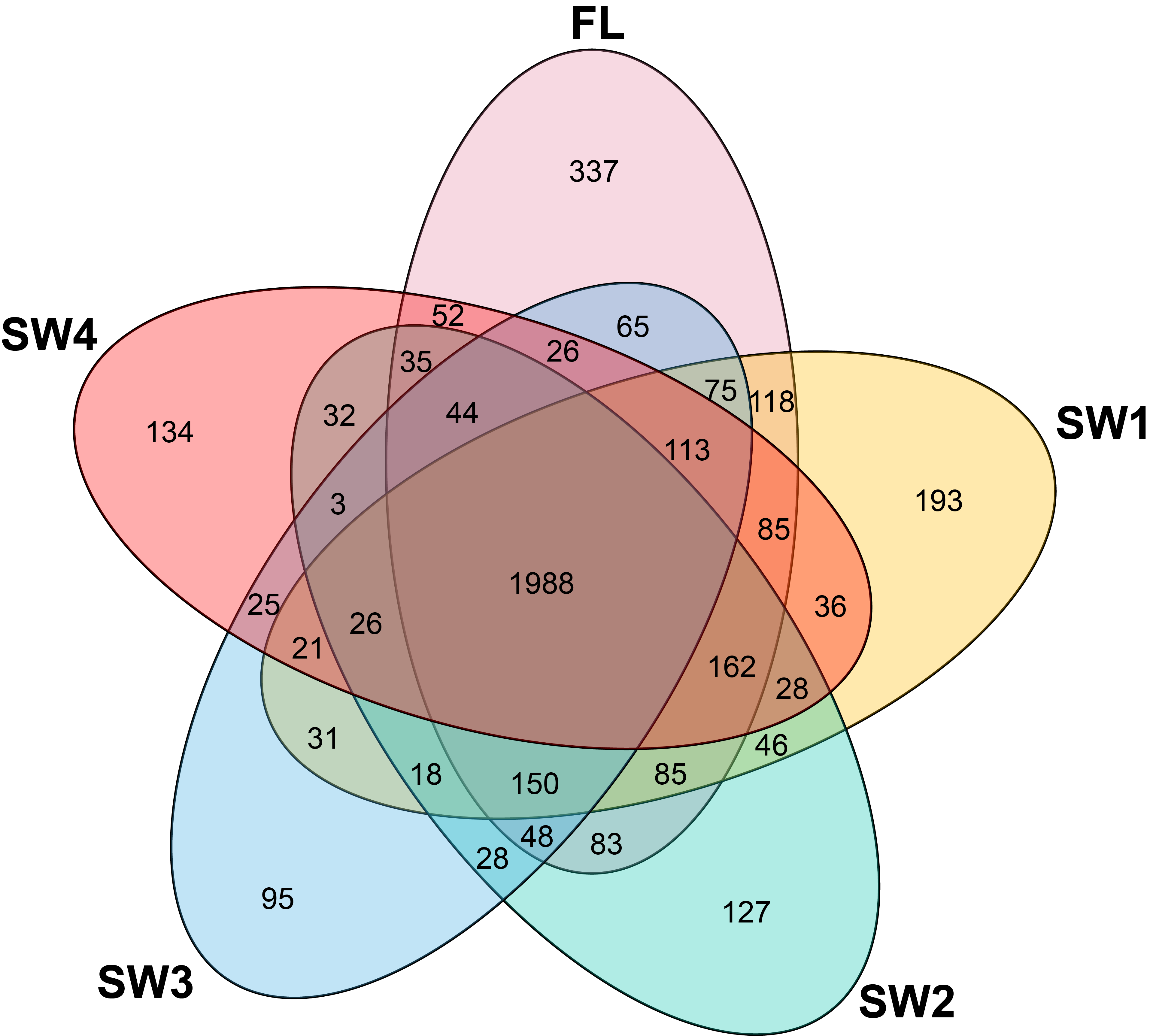

### Figure S4

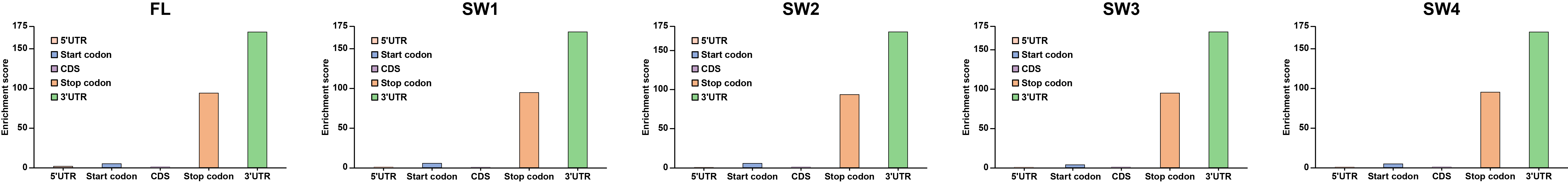

### Figure S5

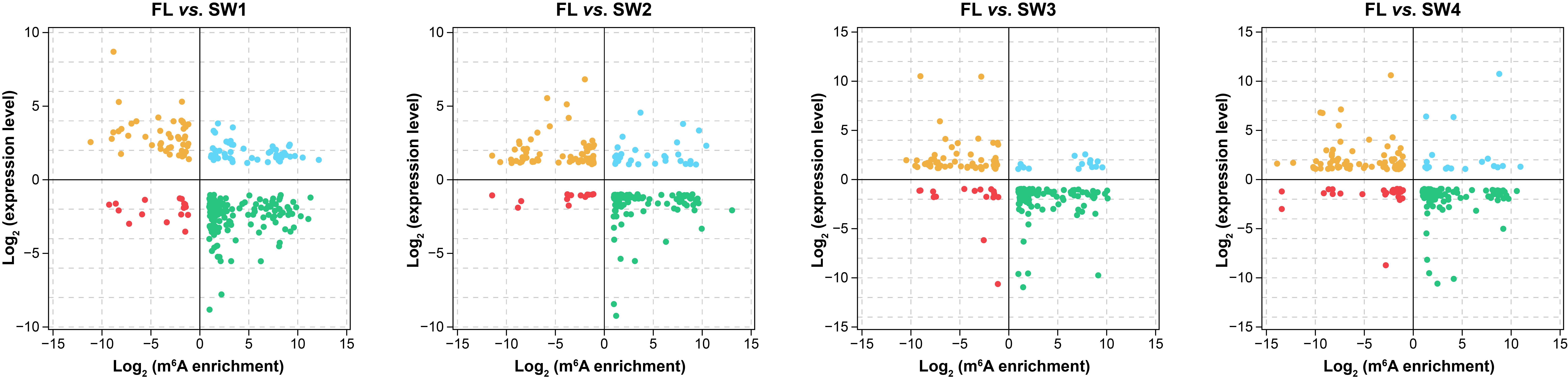

### Figure S6

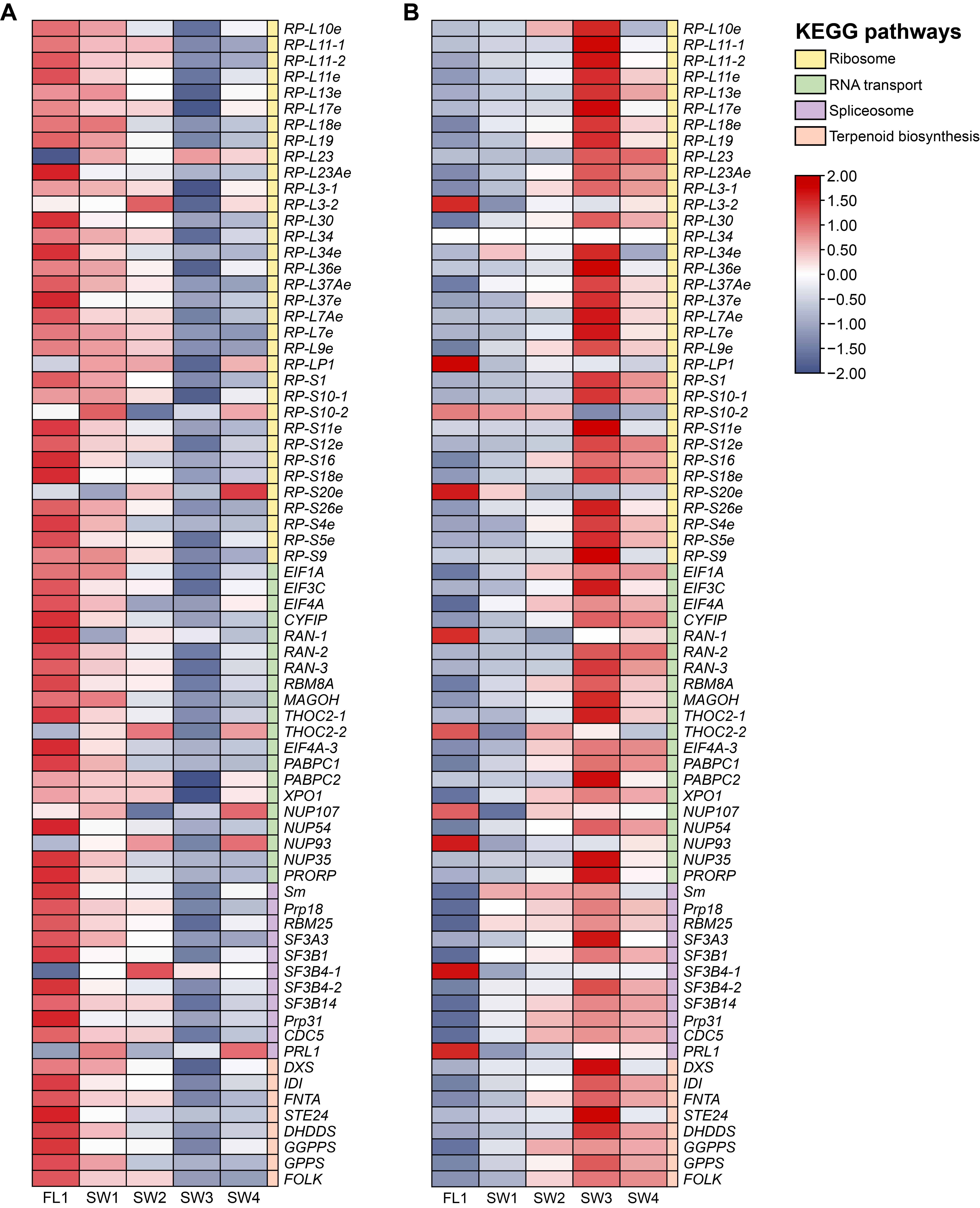

### Figure S7

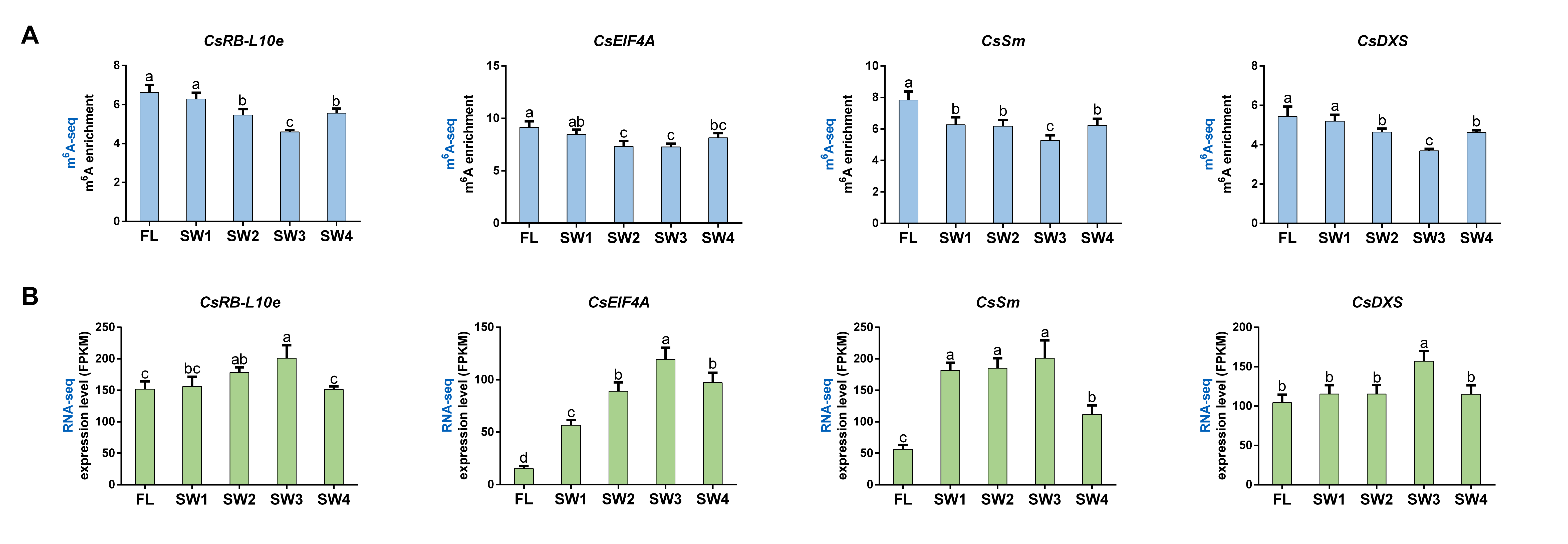

### Figure S8

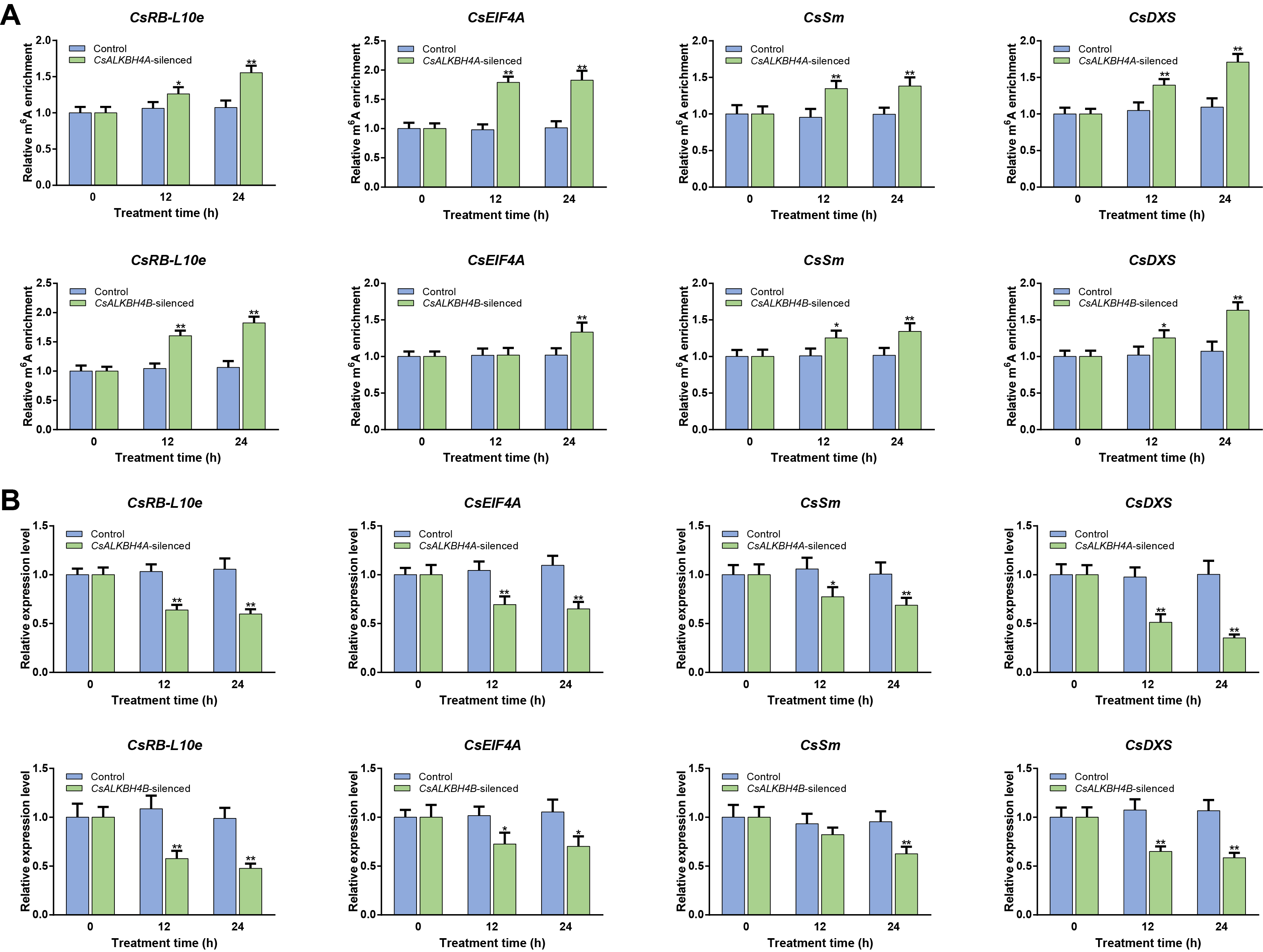

### Figure S9

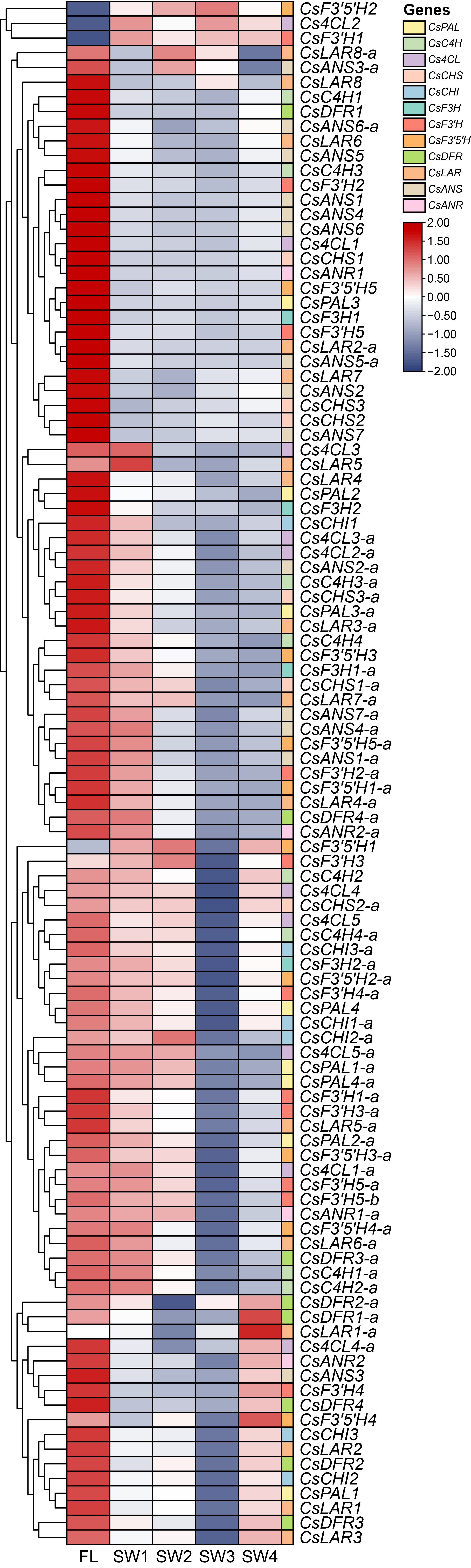

### Figure S10

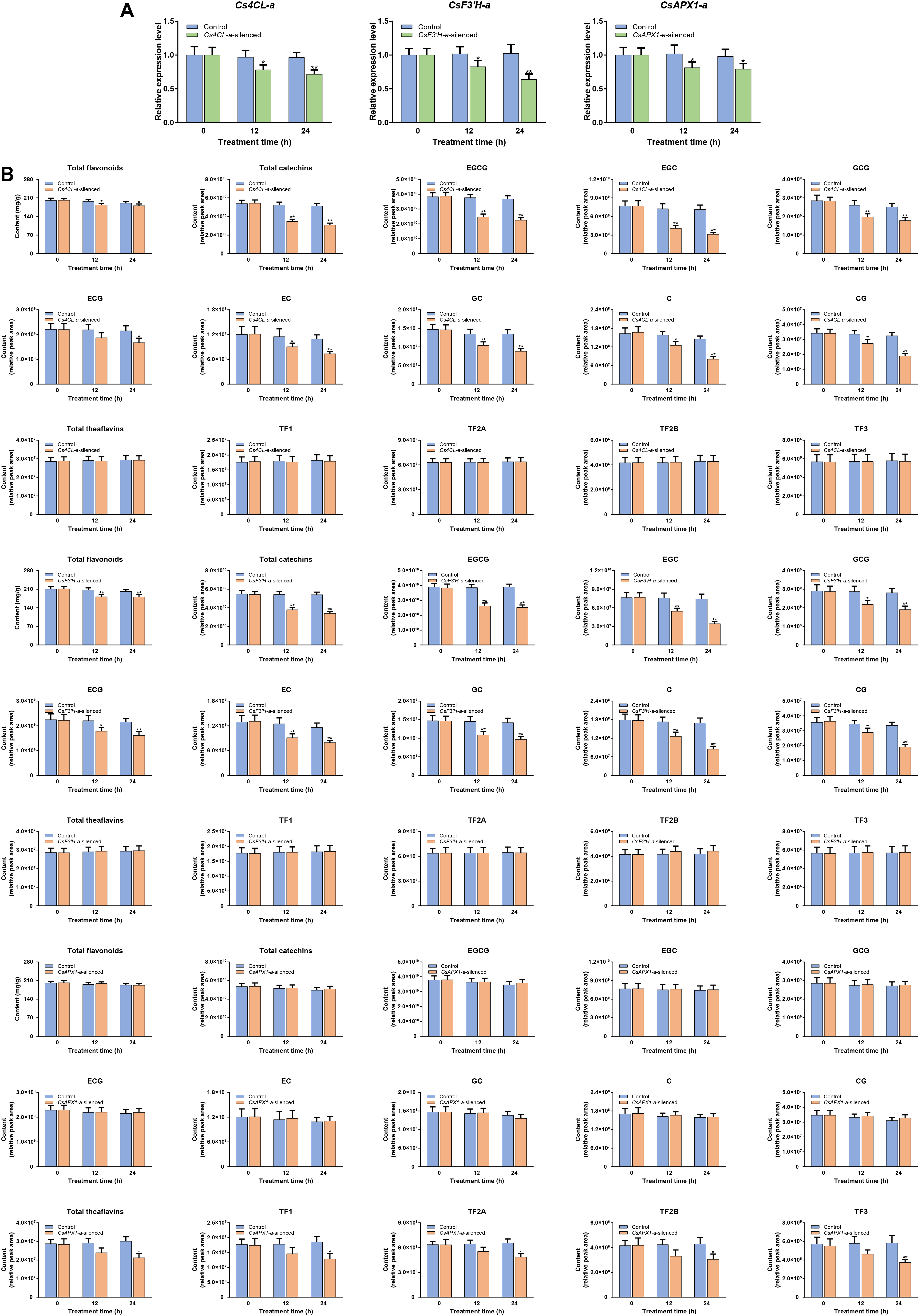

### Figure S11

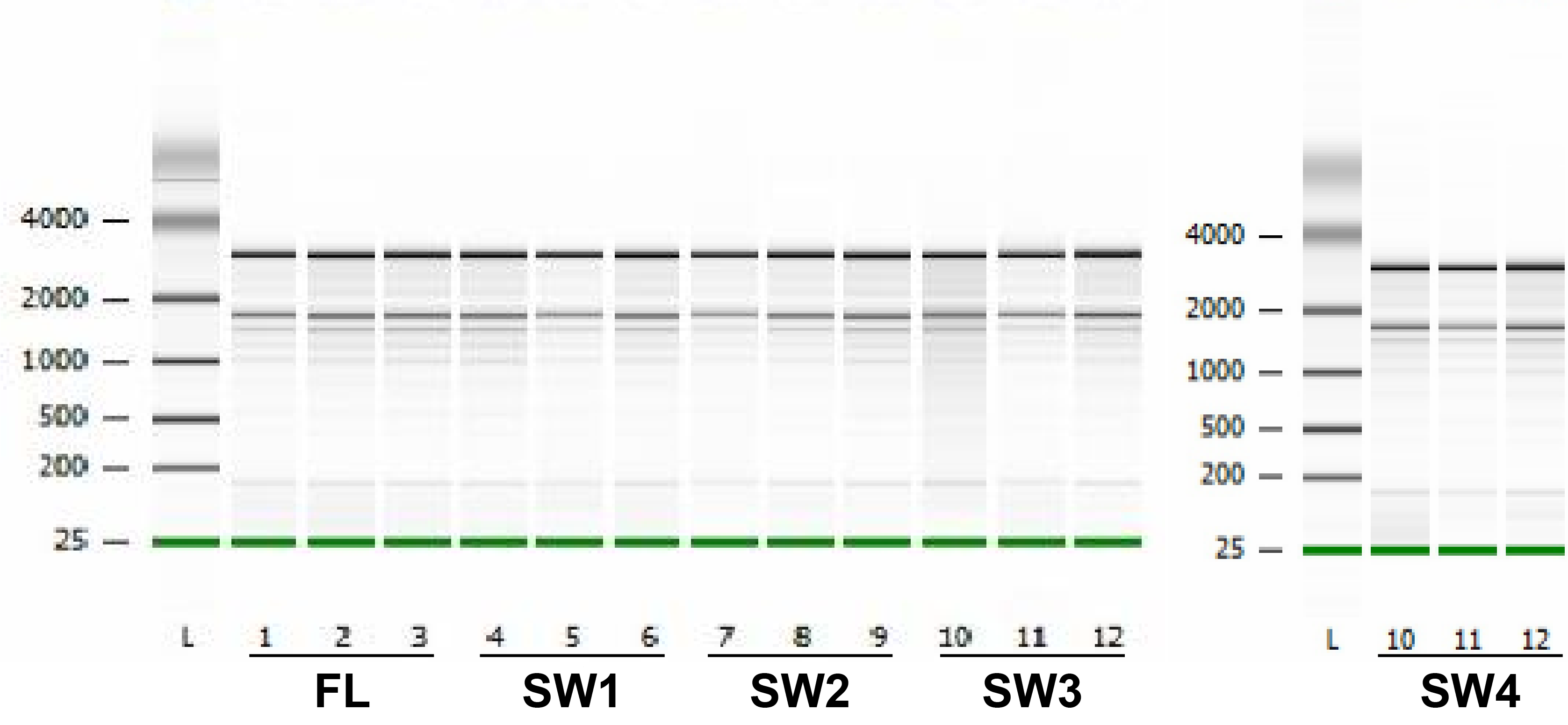
